# Cell-type specific parallel circuits in the bed nucleus of the stria terminalis and the central nucleus of the amygdala of the mouse

**DOI:** 10.1101/238493

**Authors:** Jiahao Ye, Pierre Veinante

**Author notes:** **Corresponding author**: Pierre Veinante, Institut des Neurosciences Cellulaires et Intégratives, CNRS UPR3212, 5 rue Blaise Pascal, 67084, Strasbourg, France. **Abbreviations** ac: anterior commissure ASt: amygdalostriatal transition area BDA: biotin dextran amine, 10000 MW BL: basolateral nucleus of the amygdala BLA: basolateral nucleus of the amygdala, anterior BLP: basolateral nucleus of the amygdala, posterior BMP: basomedial nucleus of the amygdala, posterior Calcrl: calcitonin receptor-like CARD: combined catalyzed reporter deposition CeA: central nucleus of the amygdala CeC: central nucleus of the amygdala, capsular part CeL: central nucleus of the amygdala, lateral part CeL/C: central nucleus of the amygdala, lateral and capsular part CeM: central nucleus of the amygdala, medial part CGRP: calcitonin gene-related peptide CGRPR: calcitonin gene-related peptide receptor CPu: caudate putamen CRF: corticotropin-releasing factor cst: commissural stria terminalis CTb: cholera toxin B subunit D2R: dopamine receptor D2 DAPI: 4’,6-Diamidino-2-Phenylindole, Dihydrochloride DMPAG: dorsomedial periaqueductal gray DR: dorsal rahpe nucleus EAc: central extended amygdala ENK: enkephalin FG: Fluorogold Fu: fusiform nucleus GI/DI: granular and dysgranular insular cortex terminalis GP: globus pallidus Htr2a: serotonin receptor 2a i.p.: intraperitoneal injection InsCx: insular cortex KLH: keyhole limpet hemocyanin LaVM: lateral nucleus of amygdala, ventromedial LPAG: lateral periaqueductal gray LPB: lateral parabrachial nucleus LPBE: external lateral parabrachial nucleus MPB: medial parabrachial nucleus NPY: neuropeptide Y PAG: periaqueductal gray PB: phosphate buffer PBS: phosphate-buffered saline PHA-L: *Phaseolus vulgaris* leucoagglutinin Pir: piriform cortex PKCδ: protein kinase C, delta type Ppp1r1b: phosphatase 1 regulatory subunit 1B positive Rspo2: R-spondin 2 positive s.c.: subcutaneous injection S2: secondary somatosensory cortex scp: superior cerebellar peduncle SEM: standard error of the mean SOM: somatostatin ST: bed nucleus of the stria terminalis STL: lateral bed nucleus of the stria terminalis STLD: dorsal lateral bed nucleus of the stria terminalis STLP: posterior lateral bed nucleus of the stria terminalis STLV: ventral lateral bed nucleus of the stria terminalis STMA: anterior medial bed nucleus of the stria terminalis STMV: ventral medial bed nucleus of the stria terminalis VLPAG: ventral lateral periaqueductal gray.

## Abstract

The central extended amygdala (EAc) is a forebrain macrosystem which has been widely implicated in fear, anxiety and pain. Its two key structures, the lateral bed nucleus of the stria terminalis (STL) and the central nucleus of amygdala (CeA), share similar mesoscale connectivity. However, it is not known whether they also share similar cell-specific neuronal circuits. We addressed this question using tract-tracing and immunofluorescence to reveal the connectivity of two neuronal populations expressing either protein kinase C delta (PKCδ) or somatostatin (SOM). PKCδ and SOM are expressed predominantly in the dorsal part of STL (STLD) and in the lateral/capsular parts of CeA (CeL/C). We found that, in both STLD and CeL/C, PKCδ+ cells are the main recipient of extra-EAc inputs from the external lateral part of the parabrachial nucleus (LPBE), while SOM+ cells are the sources of long-range projections to extra-EAc targets, including LPBE and periaqueductal gray. PKCδ+ cells can also integrate inputs from posterior basolateral nucleus of amygdala or insular cortex. Within EAc, PKCδ+, but not SOM+ neurons, serve as the major source of projections to the ventral part of STL and to the medial part of CeA. However, both cell types mediate interconnections between STLD and CeL/C. These results unveil the pivotal positions of PKCδ and SOM neurons in organizing parallel cell-specific neuronal circuits of CeA and STL, which further support the idea of EAc as a structural and functional macrostructure.

## INTRODUCTION

The central extended amygdala (EAc) is a forebrain macrosystem which contributes to diverse functions and disorders including pain, associative learning behaviors and emotion in animal models (Neugebauer et al. 2004; Shackman and Fox 2016; Veinante et al. 2013; Alheid 2003; de Olmos and Heimer 1999). The concept of EAc is also increasingly gaining importance as a fundamental structure underlying psychiatric disorders such anxiety and post-traumatic stress syndrome in human (Shackman and Fox 2016), but the organization of its neuronal microcircuits is still elusive.

The lateral part of the bed nucleus of the stria terminalis (STL) and the central nucleus of the amygdala (CeA) form the core structures of EAc, and are connected by corridor of sublenticular cells along the ventral amygdalofugal pathway and the stria terminalis (Cassell et al. 1999). In both STL and CeA, multiple subdivisions exist but different nomenclatures have been used (McDonald 1982; Sun and Cassell 1993; Chieng et al. 2006).In the rodent brain, CeA has been divided into capsular (CeC), lateral (CeL) and medial divisions (CeM) (Cassell et al. 1999; Paxinos and Franklin 2012). In mice, however, the border between CeC and CeL is more elusive that in rats and different delineations have been applied in different studies (Haubensak et al. 2010; Li et al. 2013; Kim et al. 2017). Thus we will refer to them collectively as capsular and lateral CeA (CeL/C). On the other hand, the delineation of STL subdivisions is much less consensual (Alheid 2003; Dong et al. 2001a; Gungor and Pare 2016). In this study, we divided the middle STL level into dorsal (STLD), ventral (STLV) and posterior (STLP) parts, according to Franklin and Paxinos’s mouse brain atlas (Paxinos and Franklin 2012). CeA and STL display striking similarities in cytoarchitecture, neurochemistery and connectivity (Alheid 2003; Sun and Cassell 1993). For example, both STL and CeA are targeted at mesoscopic level by similar cortical, intraamygdaloid, thalamic and brainstem afferents, and they both project to the same hypothalamic and brainstem targets (McDonald et al. 1999; Alheid 2003; Davis and Shi 1999). In addition, STL and CeA are strongly linked by subdivisions-specific interconnections and a directional bias in intrinsic EAc connections has been suggested from STLD and CeL/C to ventral STL (STLV) and CeM (Sun et al. 1991; Cassell et al. 1999).

GABAergic neurons constitutes the large majority of neurons in STL and CeA, giving rise to local inhibition (Sun and Cassell 1993; Cassell et al. 1999; Hunt et al. 2017), as well as to mutual inhibitions between STL and CeA (Sun et al. 1991; Sun and Cassell 1993; Veinante and Freund-Mercier 1998) and long range projections (Moga et al. 1989; Sun and Cassell 1993). While tract-tracing and virus tracing clearly established GABAergic projections between STLD and CeL/C, as well as STLD or CeL/C to STLV/CeM (Ciocchi et al. 2010; Li et al. 2013; Cai et al. 2014), it is still unclear which cell populations mediate such interactions. It is indeed well known that both STL and CeA contain multiple neuronal populations expressing different neuropeptides, such as somatostatin, corticotropin-releasing factor (CRF), neurotensin, enkephalin (Cassell et al. 1999; Li et al. 2013; Haubensak et al. 2010; Veinante et al. 1997), which pose a good challenge to dissect cell-type specific circuits in EAc.

Recent researches on mouse CeL/C revealed the existence of two non-overlapping neuronal groups expressing either protein kinase C delta type (PKCδ) or somatostatin (SOM), which together constitutes the majority of local GABAergic neurons (Haubensak et al. 2010). PKCδ+ and SOM+ neurons can form discreet disinhibitory circuits controlling fear learning (Ciocchi et al. 2010; Haubensak et al. 2010; Li et al. 2013; Fadok et al. 2017), anxiety (Botta et al. 2015), active defense (Yu et al. 2016), and feeding behavior (Cai et al. 2014; Campos et al. 2016). On the other hand, STL is also involved in fear response (Davis et al. 2009; De Bundel et al. 2016) and anxiety (Kim et al. 2013; Jennings et al. 2013; Mazzone et al. 2016), yet it is not clear whether STL shares some features of the cell-type specific circuits in CeA. Moreover, the involvement of PKCδ+ and SOM+ cells in projections from CeA to STL is unknown. Based on similar enrichment of PKCδ+ and SOM+ neuronal populations in STL and CeA (Lein and et al. 2007) and the idea that symmetric components of EAc can share similar organization, we hypothesize that, similar to CeA, microcircuits based on PKCδ+ or SOM+ neurons might also exist in STL and also contribute to intra-EAc circuitry.

Thus, in this study, we combined tract-tracing and immunofluorescence in mice to address the neuronal microcircuits of STL and CeA at three levels: long-range inputs, intrinsic EAc interconnectivity, and long-range outputs. Our results show that both PKCδ+ and SOM+ neuronal populations are involved in microcircuits similarly organized in CeL/C and STLD. In both CeL/C and STLD, PKCδ+ neurons are preferentially innervated by calcitonin gene-related peptide (CGRP)-positive inputs from the external lateral part of the parabrachial nucleus (LPBE), and can also integrate other long-range excitatory inputs, from insular cortex (InsCx) and posterior basolateral amygdala (BLP). This PKCδ+ population also provides the main inhibition within EAc, by projecting to CeM and STLV. On the other hand, mutual connections between STLD and CeL/C can be mediated by both cell-types. In comparison, SOM+ neurons provide the main outputs from STLD and CeL/C to extra-EAc targets, including LPBE and periaqueductal gray (PAG).

## MATERIALS& METHODS

### Animals

Adult male C57BL/6J mice of 11-12 weeks old (Charles River®, L’Arbresle, France) were housed in standard housing cages, with ad libitum access to food and water (12/12-hour light/dark cycle). In total, 27 mice were used for this study. All the experimental procedures were carried out in accordance with the regulations from European Communities Council Directive and approved by the local ethical committee (CREMEAS under reference AL/61/68/02/13).

### Stereotaxic tract-tracing

Individual animal was anesthetized by an intraperitoneal injection (i.p.) of a mixture of ketamine (87 mg/kg) and xylazine solution (13 mg/kg). Then the deep-anesthetized animal was treated with metacam (2 mg/kg, subcutaneous, or s.c.) to alleviate inflammatory response and bupivacaine (2 mg/kg, s.c.) was infiltrated on the scalp to induce local analgesia. After that, the mouse was mounted into a stereotaxic frame (Model 900, David Kopf Instrument) and a small craniotomy was made with surgical drill allowing for passage of glass pipette.

Solution of tracers were loaded into a glass pipette (tip diameter 15-25 μm) that was pulled with a P-97 micropipette puller (Sutter Instrument) and positioned according to the stereotaxic coordinates (Paxinos and Franklin 2012) (Table 1). The tracers were injected either by iontophoresis with a constant current source (Midgard Model 51595, Stoelting Co.) or by pressure injection (Picospritzer® III, Parker Hannifin Corp). Two different tracers were used for anterograde tracing. Biotin dextran amine, 10000 MW (BDA; 2% or 4% in phosphate buffer saline, PBS; cat. #D1956, Molecular Probe®) or *Phaseolus vulgaris* leucoagglutinin (PHA-L; 2.5% in phosphate buffer, PB; cat. #L-1110, Vector Laboratories®) were injected for 10-15 min (+3-5 μA, 7 s ON/OFF cycle). Three different tracers were used for retrograde tracing. First, hydroxystilbamidine methanesulfonate (cat. #A22850, Molecular Probes®) or aminostilbamidine (cat. #FP-T8135A, Interchim®) (indicated together as Fluorogold, or FG; 2% in 0.9% NaCl), was injected for 10 min (+2 μA, 3 s ON/OFF cycle). Secondly, cholera toxin B subunit (CTb; 0.25% in 0.1 M Tris buffer and 0.1% NaCl; cat. #C9903, Simga®) was injected for 15 min (+4-5 μA, 3 s ON/OFF cycle). The third tracer, red Retrobeads™ (50 -150 nl; Lumafluor Inc.) was injected into regions of interest with a Picrospritzer® III.

**Table 1.** Stereotaxic coordinates used in this study.

| **Areas** | **Coordinates** |  |  |
| --- | --- | --- | --- |
|  | **AP (mm)** | **ML (mm)** | **DV (mm)** |
| STLD | +0.20 | +0.90 | -3.30 |
| STLV | +0.20 | +0.90 | -4.00 |
| CeL/C | -1.43 | +2.35 | -3.75 |
| CeM | -1.07 | +2.20 | -4.00 |
| AI/DI | -0.23 | +3.80 | -2.10 |
| BLP | -2.45 | +3.30 | -3.90 |
| PAG | -4.47 | +0.40 | -2.70 |
| LPBE | -5.19 | +1.60 | -3.60 |

Abbreviations: AP, Anterior – Posterior axis; ML, Medial - Lateral axis; DV, dorsal - ventral axis. The stereotaxic coordinates are taken from Paxinos and Franklin's mouse brain atlas (Paxinos and Franklin 2012), with the bregma point as the origin for AP and ML axis. The DV distance was referred to its cortical surface at the corresponding AP, ML location.

After the injection, the pipette was kept in place for 5 - 10 min before withdrawing. The scalp was then closed and a lidocaine spray (2%, Xylovet®) was infiltrated near the wound. The animal was monitored by the experimenter until waking up and was placed in his home cage in the animal facility for 7 to 14 days to allow transport of the tracers.

### Sections preparation

The animal was euthanized by a lethal dose of pentobarbital (273 mg/kg, i.p.) or Dolethal (300 mg/kg, i.p.). After checking the disappearance of toe-pinch reflex, the animal was transcardially perfused with ice-cold phosphate buffer for 1 min (PB; 0.1 M, pH 7.4; 10 ml) and then with fixative (2% paraformaldehyde, in 0.1 M PB, pH 7.4; 150 ml) for 15 min. The brain was removed and placed overnight for post-fixation in the fixative (4 °C). Then, brains were kept in phosphate-buffered saline (PBS, Cat. # ET300-A, Euromedex, France; 4 °C) for one week or in PBS-sodium azide (0.02%) for longer time before sectioning. Serial coronal sections (thickness 30 μm) were cut with a vibratome (VT1000S, Leica Biosystem). Sections were kept in PBS (4 °C) for use within one week or in sodium azide (0.02% in PBS) for longer time. Subsequent immuno- and histo-fluorescence procedures were then carried out on selected brain sections (120 μm apart for adjacent slices) to for each animal. The procedures were carried out to simultaneously visualize PKCδ+ and/or SOM+ neurons together with the tracers and/or another cellular marker of interest (i.e. CGRP), through different combinations of primary and secondary antibodies.

### Combined catalyzed reporter deposition (CARD) for somatostatin

In our hands, a traditional immunofluorescent staining of SOM revealed only a few cell bodies in STLD and CeL/C, probably due to the low content of neuropeptide in the soma of projection neurons. In order to get robust staining of SOM+ cell bodies in EAc, we thus applied a highly sensitive method known as the combined catalyzed reporter deposition (CARD) (Speel et al. 1997; Hunyady et al. 1996). With the catalytic power of horseradish peroxidase, the CARD method allows specific deposits of tyramide-conjugates nearby the antigen. The reaction can amplify the immunochemical signal up to a 10 to 100- fold, compared to that of general immunofluorescent staining (Hunyady et al. 1996). In this study, we use fluorochrome-conjugated tyramide (i.e. fluorescein-tyramide and Cy3-tyramide) to reveal SOM signal. All procedures were carried out in floating brain sections, at room temperature, unless specified otherwise. First, the intrinsic peroxidase activity of brain slices was inhibited by 1% H_2_O_2_ in 50% ethanol solution for 20 min. Sections were then washed with PBS (3 x 5 min), and blocked with the blocking buffer (Triton X-100 0.3% and donkey serum 5% in PBS) for 45 min. After that, sections were incubated overnight with rabbit anti-somatostatin antibodies (Table 2) in dilution buffer (Triton X-100 0.3% and donkey serum 3% in PBS). Then, the sections were washed with PBS (3 x 5 min), and incubated with the HRP-conjugated donkey anti-rabbit antibody (1:300, in dilution buffer) for 3 hours. Sections were then washed in PBS (2 x 5 min) and then in PBS-imidazole buffer (100 mM, pH 7.6; 5 min). Finally, the CARD reaction was carried out with fluorescein-tyramide or Cy3-tyramide (1:1000, a gift from Prof. Klosen, University of Strasbourg) in PBS-imidazole buffer and H_2_O_2_ (0.001%) for up to 30 min. The reaction was stopped by washing off with PBS (3 x 5 min). The same CARD procedures were also used to reveal BDA labeled axons (i.e. Fig. 4 - 5) when the signal was weak with traditional histofluorescent staining. In those cases, peroxidase was introduced by incubation of ABC-HRP system (1: 500; Cat. # PK-6100, Vector Laboratories™) for 1.5 hr (room temperature).

**Table 2.**
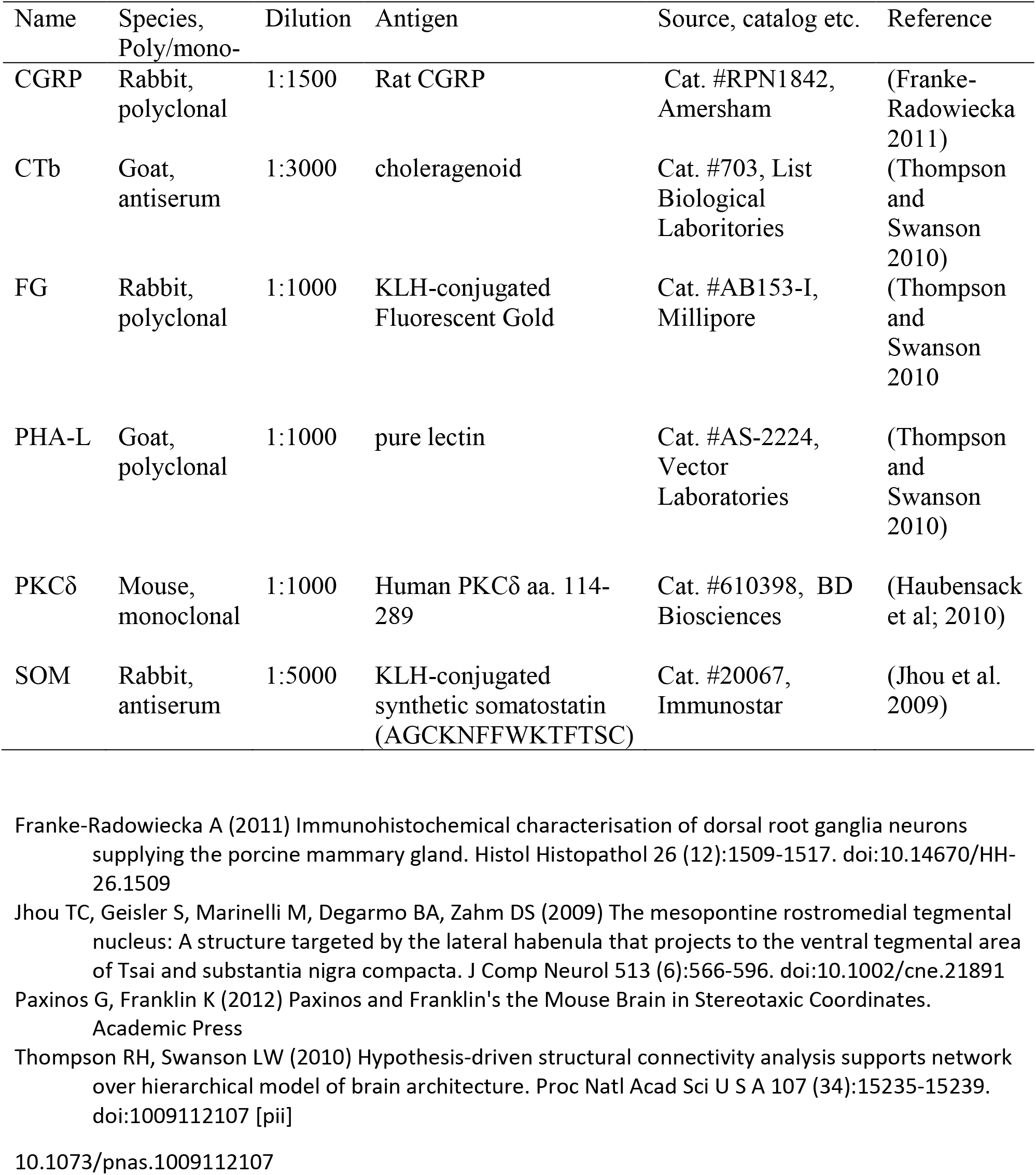
Primary antibodies

### Immunofluorescent staining

General immunofluorescent staining of other antigens were carried out after CARD revelation of SOM, when applicable. Thus, SOM immunoreactivities, together with a tracer (i.e. CTb, FG) or other cellular marker of interest (i.e. PKCδ, CGRP), were simultaneously visualized with combinations of different primary antibodies (see Table 2) and secondary antibodies, following the general procedure descibed below.

After finishing the CARD revelation of SOM, a combination of primary antibodies were applied overnight at room temperature in dilution buffer. The combinations depended on the aim of each experiment, type of tracers and technical constraints. For example, we added PKCδ primary antibodies to show the spatial distribution of PKCδ+ and SOM+ neurons, but also used PKCδ and CGRP immunofluorescence to analyze the apposition of CGRP terminals onto labeled neurons in EAc.

Sections were then washed in PBS (3 x 5 min) and incubated with corresponding secondary antibodies (1:300 in dilution buffer) for 3 hrs at room temperature. Diverse fluorophore-conjugated secondary antibodies were chosen for triple labeling of SOM, PKCδ and the third antigen, based on the compatibility of fluorophores. Overall, the following secondary antibodies were used: donkey anti-mouse-Alexa-647 conjugates (Cat. #: A-31571, Invitrogen™), donkey anti-mouse-Cy3 conjugates (Cat. #: 715-165-151, Jackson Immunoresearch™), donkey anti-rabbit-Cy5 (Cat. #: 711-175-152, Jackson Immunoresearch™), donkey anti-rabbit-Alexa 488 (Cat. #: A-21206, Invitrogen™), donkey anti-goat-Alexa 488 (Cat. #: A-11055, Invitrogen™). Streptavidin-Alexa 488 conjugate (1: 750; Cat. #: S32354, Molecular Probe®) was used for the visualization of BDA.

After washing in PBS (3 x 5 min), the sections were counterstained with DAPI (4’,6-Diamidino-2-Phenylindole, Dihydrochloride; 300 nM, Cat.# D1306, Invitrogen™) for 3 - 5 min. The sections were then arranged onto Superfrost® plus slides (Thermo Fisher Scientific™) and mounted in Fluoromount™ medium (Cat. #: F4680, Sigma-Aldrich™).

The use of CARD revelation made it possible to stain two differnet of antigens with two different primary antibodies from the same species (Hunyady et al. 1996). In this study, we used different rabbit antibodies for SOM, FG, and CGRP. For instance, to simultaneously visualizing of PKCδ, SOM, and CGRP, a low concentration of rabbit-anti-SOM (1: 5000) was used for CARD revelation, and a higher concentration of rabbit-anti-CGRP (1: 1000) antibody was subsequently applied. In this way, SOM and FG or CGRP could be revealed with sequential applications of primary antibodies from rabbit, without showing detectable cross-staining. The absence of cross-staining was determined by the separation of the staining pattern and negative control experiments in which CGRP primary antibodies were omitted.

### Imaging and analysis

For each animal, the location of injection core of tracer was examined on successive sections containing the injection sites and was evaluated according to salient anatomical features (i.e. fiber bundles) and neurochemical features (i.e. DAPI staining, PKCδ+ immunoreactivity). The delineation of subdivisions of EAc, LPB, PAG, INsCx and BLP weas done according to the fourth edition of mouse brain atlas (Paxinos and Franklin 2012). Cases in which the injection sites spilled beyond the target in nearby regions were not included into the data analysis.

For illustrations of injection sites and neurochemical patterns, if not stated otherwise, epifluorescence images were acquired by an Axio Imager 2 (Carl Zeiss™) microscope equipped with a digital camera (ProgRes® CF*cool,* Jenoptik, GmbH, Germany), under 10x, or 20x objectives; or by a NanoZoomer S60 (Hamamatsu Photonics) under a 20x objective.

To demonstrate the co-localization of markers and potential appositions between neurons and axonal processes, confocal imaging at the middle focal plane of the section was taken with a Leica TCS SP5 II system (Leica Biosystem). Images were sampled to pixel resolution = 0.255 μm by 2.5-fold of Nyquist sampling, under 20x objective with 1 airy unit. To gain more details of axonal apposition, single plane or z-stack (1 μm) confocal images were taken under 63x objective. For quantitative analysis of colocalization, epifluorescent images were taken with Axio Imager 2, under 20x apochromatic objectives. A z-stack image (step size = 2.049 μm) was obtained in STLD (bregma +0.13 mm) or CeA (bregma -1.43 mm) for each animal. The colocalization of tracers with PKCδ+ or SOM+ neurons in epifluorescence was also confirmed by corresponding confocal images. Preprocessing of images, primarily for pseudo-coloring and adjusting contrast, and subsequent analysis including cell counting and colocalization were carried out manually on open software FIJI (Schindelin et al. 2012).

### Statistics

For colocalization and apposition studies, mean value and standard error of the mean (SEM) are reported by injection group and brain areas. Unpaired two-sample Student’s *t*-test was carried out in R program (©The R Foundation).

## RESULTS

### Distribution of PKCδ neurons and SOM neurons in STLD and CeL/C

We first examined the pattern of PKCδ and SOM imunoreactivities in subdivisions of the STL (n = 3) and the CeA (n = 3). PKCδ+ soma were detected mainly in the STL and CeA, as well as in the lateral septum (Fig. 1a), the thalamus (Fig. 1d). In STL, well-stained PKCδ+ cell bodies were concentrated in the STLD of which they sharply defined its limits with surrounding STLP (Fig. 1a). In CeA, PKCδ+ soma were present in CeL/C where they tend to be concentrated laterally with a reduced density medially, at the limit with the CeM (Fig. 1d). Dense PKC δ+ neuropil was also obviously packed in STLD and CeL/C (Fig. 1a, d; see also Fig. 2c, e, f). SOM+ neurons were observed mainly in the STL, cerebral cortex, caudate-putamen, hypothalamus (Fig. 1b), and amygdala (Fig. 1e). While the staining of SOM+ interneurons in cerebral cortex and caudate-putamen filled the cell bodies, the SOM labeling of somas in STL and CeA was patchier and hardly defined the somatic contour, probably due to the low content of SOM in the soma of projection neurons. In the STL, SOM+ neurons and fibers were observed in all subdivisions, but appeared denser in the STLD (Fig. 1b), where their distribution overlapped with that of PKCδ+ neurons (Fig. 1c). In the CeA, a low density of SOM+ soma and processes appeared in the CeM, but a strong concentration was observed in the CeL/C (Fig. 1e). The distribution of SOM+ cell bodies overlapped with that of PKCδ+ neurons medially (i.e. CeL), but decreased laterally (i.e. CeC), where PKCδ+ neurons were the most abundant (Fig. 1f). Despite their similar regional distribution in STLD and CeL/C, PKCδ and SOM immunoreactivities remained segregated at cellular level and were almost never observed in the same neurons (Fig. 1c, f; see also Fig2c, e). Finally, while PKCδ+ and SOM+ neurons were observed along the rostrocaudal extent of STLD (bregma +0.25 mm to +0.01 mm) and CeL/C (bregma -0.80 mm to -2.03 mm), their density appeared stronger in the caudal parts of STLD and CeL/C.

**Fig 1.**
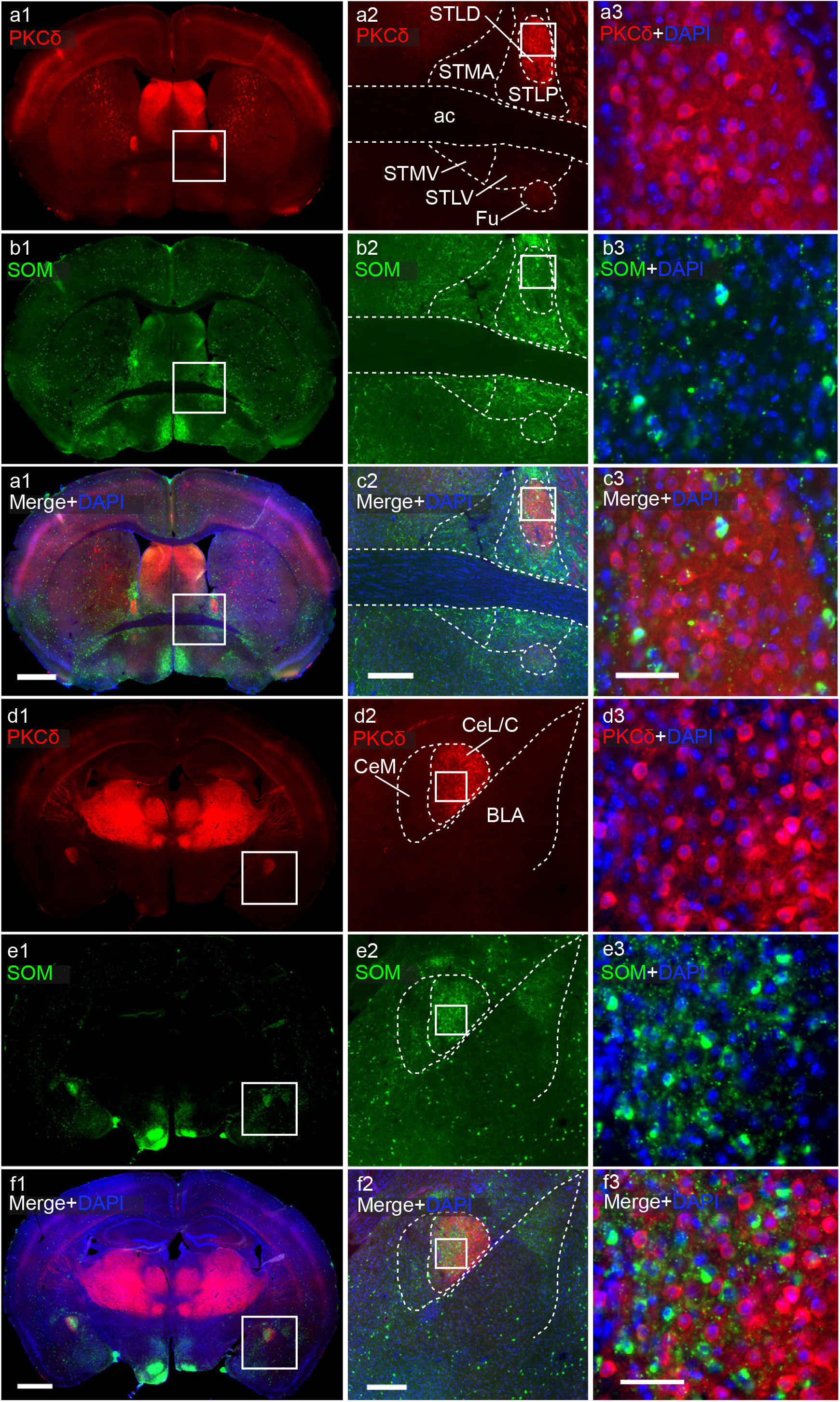
PKCδ and SOM expressing cells are concentrated in STLD and CeL/C. Double staining of PKCδ (a, c, d, f; red) and SOM (b, c, e, f; green) in coronal sections of STLD (a - c; bregma level +0.13mm) and CeL/C (d - f; bregma -1.55 mm), detected with epifluorescence. DAPI staining (blue) of cell nuclei is also shown (c1, f1, c2, f2 and a3 - f3). The first column shows a view of full sections at the level of the STL (al - cl) and of the amygdala (d1 – f1); the second column shows a detailed view of STL (a2 - c2) and amygdala (d2 - f2) with delineations, corresponding to the boxed area in al-fl; the third column shows a magnification at cellular level in the STLD (a3 - c3) and in the CeL/C (d3 - f3) of the boxed areas in a2 - f2. See list for abbreviations. Scale bars: al-fl, 1.0 mm; a2 - f2, 500 μm; a3 - f3, 50 μm.

**Fig 2.**
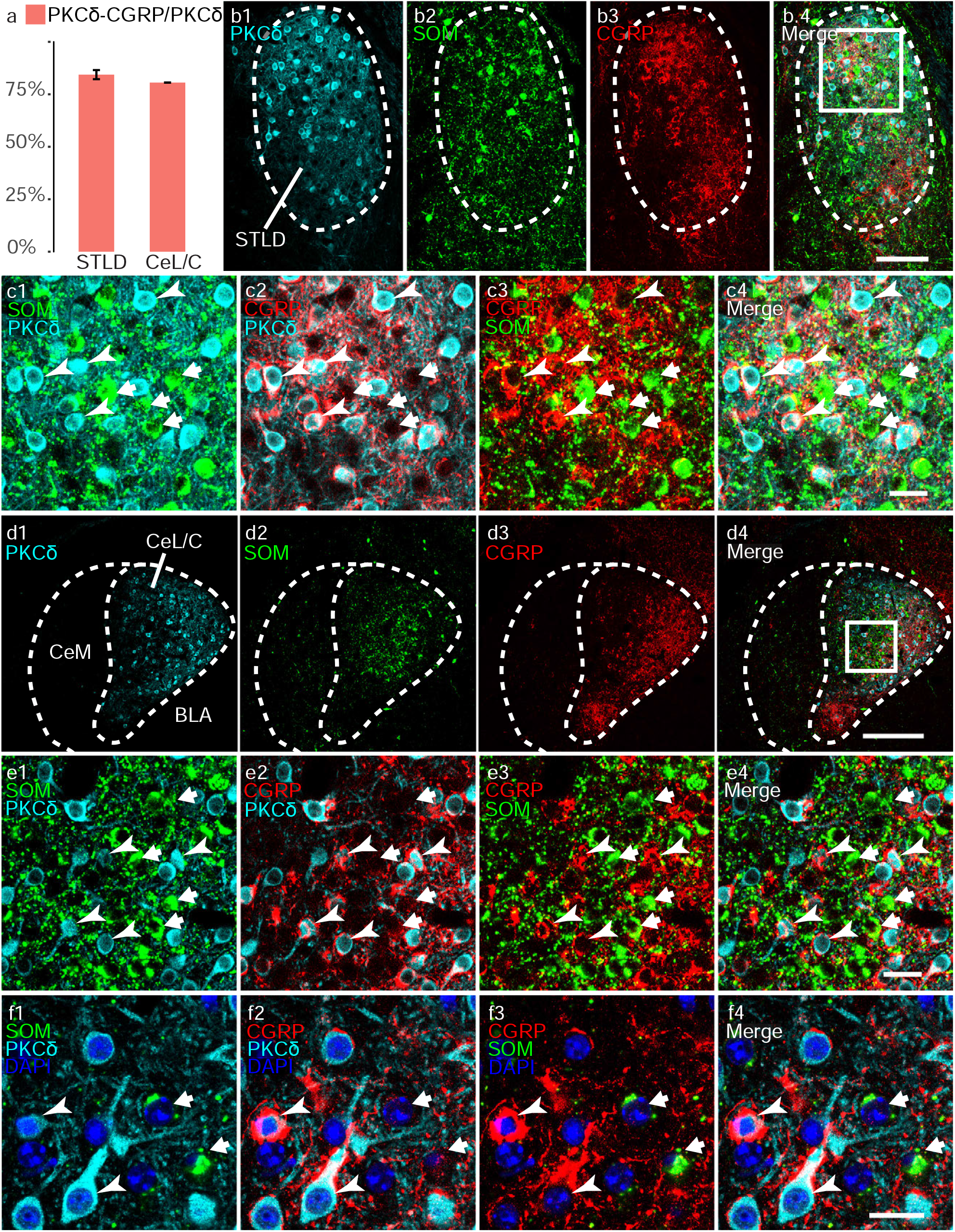
Structural apposition of CGRP+ terminals onto PKCδ+ neurons in STLD and CeL/C. Confocal imaging of triple labeling for PKCδ (cyan), SOM (green) and CGRP (red) in STLD (b, c) and CeL/C (d - f). a Percentages of PKCδ+ somas in putative contact with perisomatic CGRP+ terminals, for STLD (Mean = 84.4%, SEM = 0.031, n = 3) and CeL/C (Mean = 80.6%, SEM = 0.0005, n = 3). b1 - b4, d1 - d4: Low power view of STLD (b1 – b4) and CeL/C (d1 - d4) showing the distribution of PKCδ (b1, d1), SOM (b2, d2),CGRP (b3, d3) immunoreactivities and the three signals merged (b4, d4). c1-c4, e1-e4: Magnifications at cellular level of the boxed areas in b4 (STLD) and d4 (CeL/C) showing signals for PKCδ and SOM (cl, el), CGRP and PKCδ (c2, e2), CGRP and SOM (c3, e3) and merge (c4, e4); the arrows point to PKCδ+ neurons and the arrowheads point to SOM+ neurons (same in f1 - f4). Note the absence of overlap between PKCδ+ and SOM+ somas (c.1,e.1), the frequent appositions of CGRP+ baskets around PKCδ+ somas (c2, e2) and the absence of such appositions onto SOM+ somas (c3, e3). In f1 - f4, a further magnification in STLD leads to the same observations and shows that CGRP+ baskets wrapped around soma and primary dendrites of PKCδ+ somas. Abbreviations, see the list. Scale bars: b.1-b4, 100 μm; c1 - c4, 25 μm; d1 - d4, 200 μm; e1 - e4, 25 μm; f1 - f4, 20 μm.

Thus, we confirmed the expression of the similar cellular markers, PKCδ and SOM, in segregated neuronal populations of CeL/C, in accordance with previous descriptions (Ciocchi et al. 2010; Li et al. 2013; Haubensak et al. 2010), and showed that a similar situation occurs in STLD..

### A majority of PKCδ+ neurons are closely surrounded by CGRP+ terminals

Having established the distribution of PKCδ+ and SOM+ neurons in STLD and CeL/C, we tested whether external inputs could target similar populations in both nuclei. The lateral parabrachial nucleus (LPB) is known to provide a dense input to STLD and CeL/C (Bernard et al. 1993; Alden et al. 1994). This LPB-EAc pathway is characterized by large basket-like pericellular terminals (Sarhan et al. 2005) co-releasing glutamate and neuropeptides, especially CGRP (Delaney et al. 2007; Salio et al. 2007). As the CGRP innervation to EAc has been shown to originate essentially from LPB in rats (Yasui et al. 1991b; D’Hanis et al. 2007), and as a recent study in mice suggested that the cells expressing CGRP receptor overlap with SOM and PKCδ populations (Han et al. 2015), we first examined the potential innervation of SOM and PKCδ by CGRP terminals using a triple immunofluorescence protocol (Fig. 2).

In accordance with previous descriptions, CGRP+ terminals were observed in the STLD and the CeL/C. Their distribution largely overlapped with that of PKCδ+ cells and partially overlapped with that of SOM+ cells (Fig. 2b, d) and displayed characteristic perisomatic terminals (Fig. 2c, e).

Confocal analysis at cellular level showed that PKCδ+ somas were often surrounded by basket-like CGRP+ elements in STLD (Fig. 2c) and CeL/C (Fig. 2e). A close observation revealed the wrapping of soma and proximal dendrites of PKCδ+ neurons by CGRP+ terminals (Fig. 2f). A quantitative analysis (n = 3) indicated that 84.4% and 80.6 % of PKCδ+ soma in STLD and CeL/C, respectively, were closely surrounded by CGRP+ perisomatic terminals (Fig. 2a). In addition, most of CGRP+ baskets-like structures contact either PKCδ+ neurons or PKCδ-/SOM-neurons.

By contrast, CGRP+ basket-like structures almost never surrounded SOM+ somas in STLD (Fig. 2c) or CeL/C (Fig. 2e, f). Yet, we cannot exclude that thinner single CGRP+ terminal lacking the basket-like appearance, could contact SOM+ neurons, as such putative appositions were sometimes registered under high magnification (Fig. 2f). However, the incomplete staining of SOM+ soma did not allow validating the existence of such contacts.

Thus, these evidences support a dominant perisomatic CGRP+ innervation onto PKCδ+, but not SOM+, neurons in EAc, eventhough an underestimated number of SOM+ neurons in STLD and CeL/C were labeled in our study. In addition, non-perisomatic contacts between CGRP+ terminals and SOM+ neurons can not be excluded.

### CGRP terminals from LPB target PKCδ neurons in EAc

In order to further confirm the possibility that CGRP+ axonal terminals contacting EAc PKCδ+ neurons were derived from the LPB, we performed BDA anterograde tracing from LPB, followed by subsequent triple fluorescent labeling.

BDA injection sites in LPB (n = 5) were centered in the LPBE (mainly from bregma -5.07 mm to -5.41 mm), with occasional expansion into neighbouring central lateral and dorsal subnuclei (LPBcl, LPBd), but never extending to medial parabrachial nucleus or Kölliker-Fuse nucleus (Fig. 3a, f). In the ipsilateral EAc, BDA+ axons were primarily located in the oval-shaped STLD (Fig. 3b, g), fusiform nucleus of ventral STL (not shown), and CeL/C (Fig. 3d, i), with only a few axonal processes in STLP or CeM. At higher magnification, distinct BDA+ perisomatic arrangements were observed, along with individual fibers (Fig. 3c, e, h, j). The comparison of BDA+ and CGRP+ signals showed that a substantial number of the BDA+ axons forming basket-like structures also contained CGRP signal. Conversely, CGRP+ basket-like structures were often coincident with BDA+ labeling (Fig. 3c, e, h, j). However, some CGRP+ perisomatic formations appeared to be BDA-, and individual BDA+ axons only partially overlapped with CGRP immunoreactivity.

**Fig 3.**
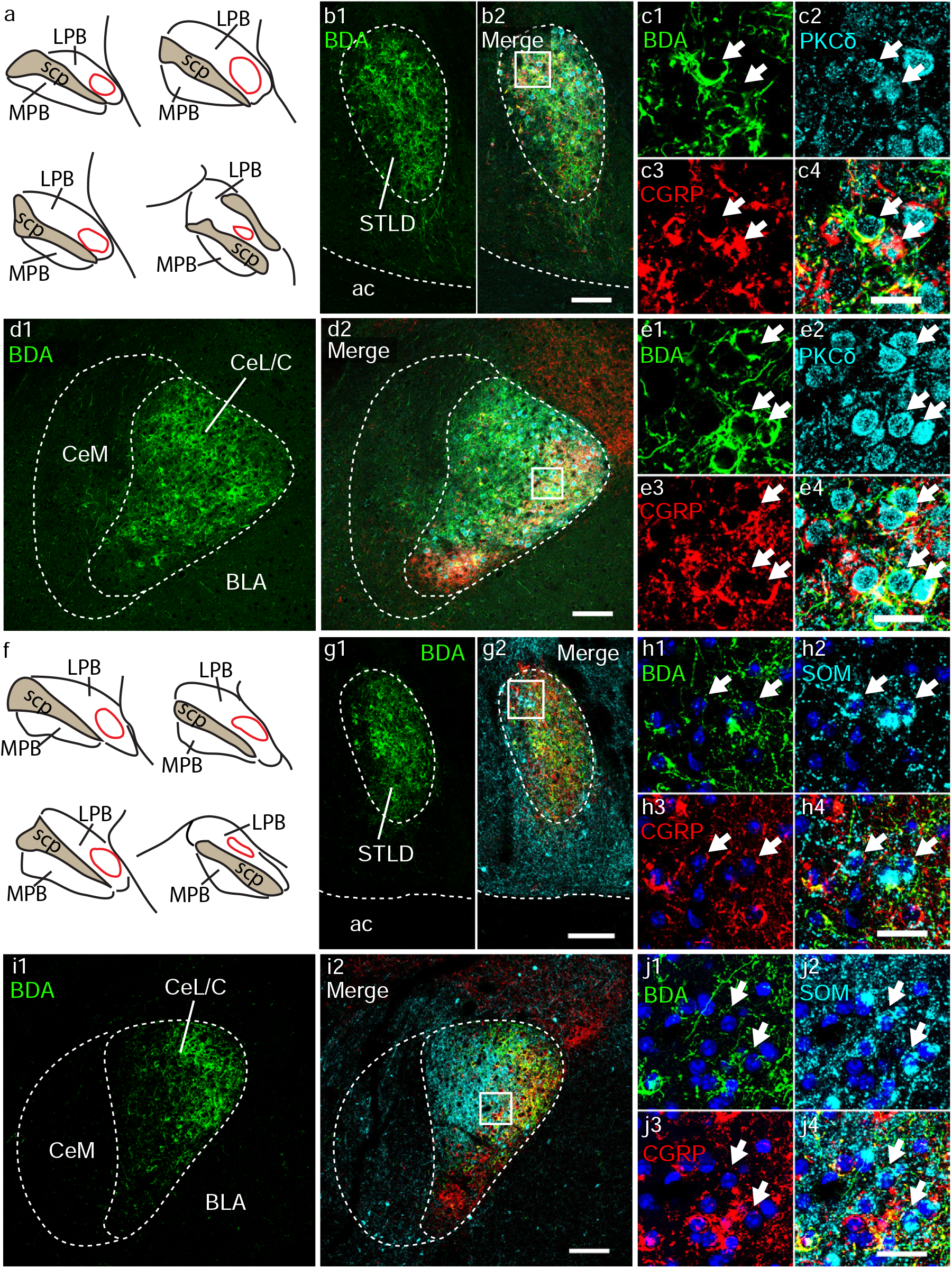
Structural apposition of CGRP+ terminals anterogradely labeled from LPB onto PKCδ+ neurons in STLD and CeL/C. Following BDA injection in the LPBE (a, f), triple labeling for BDA (green), CGRP (red) and PKCδ (cyan) (b - e) or for BDA (green), CGRP (red) SOM (cyan) along with DAPI (blue) (g - j) was performed on STLD (b, c, g, h) and CeL/C (d, e, i, j). The injection (red outlines) were centered in LPBE (a, f). Dense cores of BDA labeled fibers were observed in STLD but not STLP (b1, g1) where they overlap with the distribution of CGRP+ and PKCδ+ (b2) or SOM+ (g2) somas and fibers. Similarly, BDA-labeled fibers were densest in CeL/C (d1, i1) and partially overlapped with the distribution of CGRP+ and PKCδ+ (d2) or SOM+ (i2) somas and fibers. In c1 - c4 and e1 - e4, the higher magnifications of the boxed areas in b2 and d2, respectively, show that BDA+ basket-like structures, either in CGRP+ or CGRP-, can be found in close apposition with PKCδ+ somas (arrows) in STLD (c1 - c4) and CeL/C (e1 - e4). In addition, BDA-/CGRP+ terminals can also contact PKCδ+ somas. In h1 - h4 and j1 - j4, the higher magnifications of the boxed areas in g2 and i2, respectively, show that BDA+ basket-like structures, either CGRP+ or CGRP- are rarely found in close apposition with SOM+ somas in STLD (h1 - h4) and CeL/C (j1 - j4). Abbreviations: see list. Scale bars: b1 - b2, 100 μm; c1 - c4, 25 μm; d1 - d2, 100 μm; e1 - e4, 25 μm; g1 - g2, 100 μm; h1 - h4, 25 μm; i1 - i2, 100 μm; j1 - j4, 25 μm.

Triple labeling for PKCδ, CGRP and BDA (Fig. 3a - e) revealed that the large majority of the PKCδ+ somas in STLD (Fig. 3c) and CeL/C (Fig. 3e) were surrounded by CGRP+ baskets, as shown in the previous experiment, including most of the BDA+/CGRP+ baskets. In addition, a number of BDA+/CGRP-axonal segments were also found in close apposition with PKCδ+ somas. In sections processed for triple labeling for SOM, CGRP and BDA (Fig. 3f - j), perisomatic structures revealed by BDA and/or CGRP signals very rarely contacted SOM+ cell bodies, albeit BDA+/ CGRP-terminals could be found in close proximity to SOM+ somas in STLD (Fig. 3h) and CeL/C (Fig. 3j).

Thus, the preferential perisomatic CGRP innervation onto PKCδ+, but not SOM+, neurons in STLD and CeL/C, is likely to derive, at least in part, from the LPBE. In addition, the observation of BDA+/CGRP-perisomatic terminals surrounding PKCδ+ neurons and of individual axons found close to PKCδ+ or SOM+, suggest the existence of CGRP and non-CGRP inputs from LPBE to EAc.

### PKCδ+ neurons in EAc integrate convergent signals

Beside inputs from the LPBE, both STL and CeA are strongly innervated by the basolateral nucleus of amygdala, especially its posterior subdivision (BLP) (Dong et al. 2001a; Pitkanen et al. 2003), and by the insular cortex (InsCx) (Saper 1982; Yasui et al. 1991a; Sun et al. 1994). Kim and colleagues (Kim et al. 2017) recently showed that BLP strongly targeted PKCδ+ neurons in CeL/C, and a recent study using rabies virus tracing unveiled convergent inputs to CeL PKCδ+ neurons from multiple brain regions including InsCx, BLP and LPBE (Cai et al. 2014). However, it is not known if the same goes true for STLD PKCδ+ neurons and whether they can potentially integrate information from both intra-amygdaloid (i.e. BLP) and extra-amygdaloid (i.e. LPB or InsCx) inputs. We thus injected the anterograde tracer BDA in BLP or in InsCx and carried out triple fluorescent labeling in STLD and CeL/C to look for the potential innervation of PKCδ+ neurons by CGRP+ baskets (potentially from LPBE) and BLP or InsCx afferents.

The BDA injection sites in BLP were largely confined to the lateral part of the caudal BLP (Fig. 4a, b; bregma -2.45 mm), with minor leakage in the nearby piriform cortex and lateral nucleus of amygdala. In the ipsilateral STL, BDA+ axon terminals spread quite evenly in STLD and STLP (Fig. 4c). At higher magnification, BDA+ axonal varicosities (Fig. 4d) could be observed at close appositions with PKCδ+ neurons, simultaneously surrounded by CGRP+ terminals. Similarly, the CeA was also densely innervated by BDA+ axons from BLP (Fig. 4e). At cellular level, these BDA+ axonal varicosities in CeL/C could also form close apposition with PKCδ+ neurons contacted by CGRP+ baskets (Fig. 4f).

**Fig 4.**
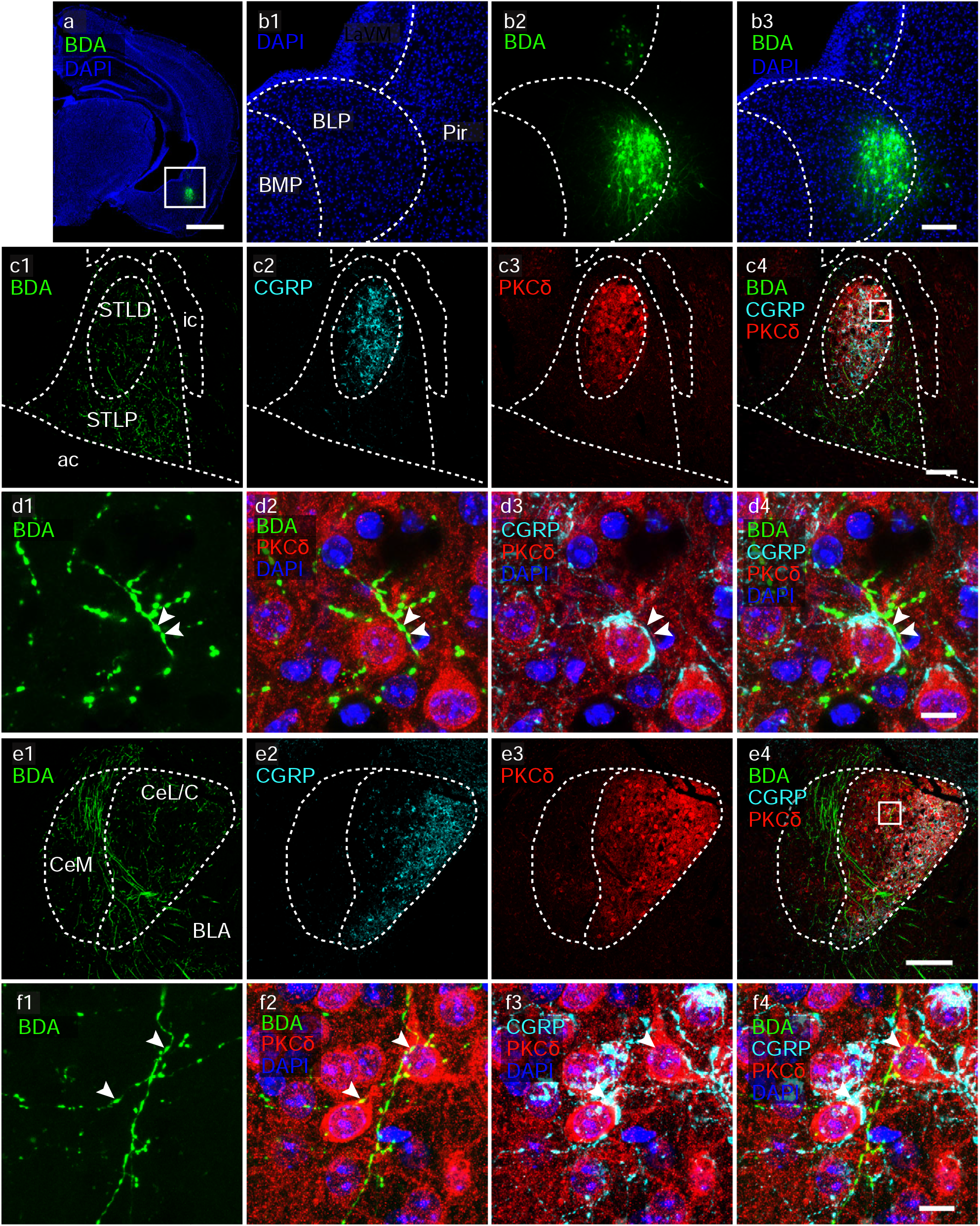
Projections from caudal BLP and from CGRP+ terminals can target the same PKCδ+ neuron in STLD and CeL/C. After anterograde tracing from the caudal BLP area (a, b; bregma level -2.45 mm), triple immunofluorescent labeling of BDA (green), CGRP (cyan) and PKCδ (red), together with nuclear counterstaining by DAPI (blue), was performed on STL (c, d) and CeA sections (e, f). BDA injections were located in lateral region of the caudal BLP, with minor leakage in the nearby piriform cortex (Pir) and ventromedial part of the lateral nucleus of amygdala (LaVM) (b1 – b3). BDA+ axon were present in most of the STL, and overlapped in STLD with PKCδ+ neurons and CGRP+ terminals. At high magnification, z-projection images (z stack = 5.43 μm) revealed close apposition of BDA+ axonal varicosities (dl, d2, d4; arrowheads) onto CGRP-inner-vated PKCδ+ neurons (d4). Similarly, moderate to dense labeling of BDA+ axonal terminals were observed in CeL/C, overlapping with CGRP+ axonal field and PKCδ+ neuronal populations. z-projection images (z stack = 9.38 μm) showed close apposition of BDA+ axonal varicosities (fl, f2, f4; arrow heads) onto PKCδ+ neurons innervated by CGRP+ axonal terminals. Abbreviations: see list. Scale bars: a, 1000 μm; b, 150 μm; cl – c4, 100 μm; dl – d4, 10 μm; el – e4, 150 μm; fl – f4, 10 μm.

The BDA injections in InsCx targeted the granular and dysgranular insular areas at middle level (bregma -0.23 mm), with some minimal extent dorsally in the secondary somatosensory cortex (S2) (Fig. 5a, b). Ipsilaterally, a moderate to strong projection was found in the STLD (Fig. 5c) and in the CeL/C (Fig. 5e), in the regions where intense CGRP+ axonal field and PKCδ+ neurons coexisted. Observation at high magnification confirmed the existences of simultaneous axonal appositions by BDA+ varicosities and CGRP+ varicosities onto a single PKCδ+ neuron in STLD (Fig. 5d) and in CeL/C (Fig. 5f).

**Fig 5.**
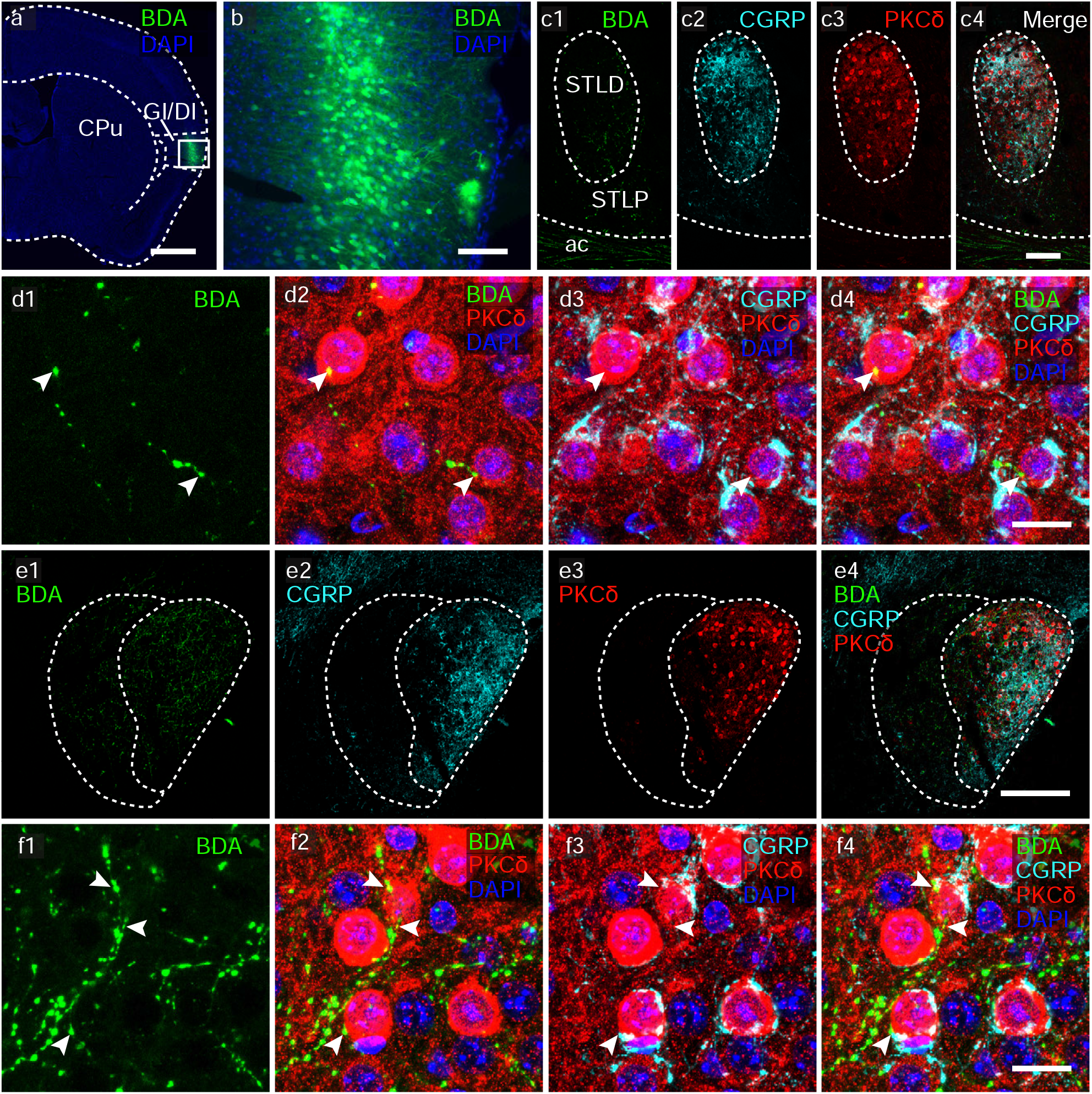
Projections from InsCx and from CGRP+ terminals can target the same PKCδ+ neuron in STLD and CeL/C. Following anterograde tracing from InsCx (a, b; bregma level -0.23 mm mm), triple immunofluorescent labeling of BDA (green), CGRP (cyan) and PKCδ (red), together with nuclear counterstaining by DAPI (blue), was performed on STL (c, d) and CeA sections (e, f). BDA injections were restricted to in layer II/III of InsCx and largely confined to granular (GI) and dysgranular (DI) areas (a, b; epifluorescent images by NanoZoomer S60). The BDA+ axons spread in all the STL, including STLD where it overlapped with PKCδ+ neurons and CGRP+ terminals (c1 - c4; single confocal plane). With high magnification, z-projection images (z stack = 8.89 μm) revealed close apposition of BDA+ axonal varicosities (d1, d2, d4; arrow heads) onto CGRP-innervated PKCδ+ neurons (d4). Similarly, BDA+ axonal terminals were also found in CeL/C, which again largely coincides with CGRP+ axonal field and PKCδ+ neuronal populations (el – e4; z stack = 5.93 μm). Higher magnification revealed close apposition of BDA+ axonal varicosities (arrow heads) onto PKCδ+ neurons surrounded by CGRP+ basket-like terminals (fl, f2, f4; z stack = 9.38 μm). Abbreviations: see list. Scale bars: a, 1000 μm; b, 100 μm; cl – c4, 100 μm; dl – d4, 15 μm; el – e4, 200 μm; fl – f4, 15 μm.

Thus, these structural evidences support the notion that PKCδ+ neurons in EAc can mediate the integration of both viscero- and somato-sensory signals from LPBE and highly processed polymodal information from BLP and InsCx. However, it should be noted that these BLP and InsCx inputs onto PKCδ+ neurons are not exclusive, as numerous BDA+ varicosities were observed without evident apposition to PKCδ+ neurons in STLD and CeL/C.

### A majority of CeM-projecting or STLV-projecting neurons in STLD and CeL/C express PKCδ

After establishing the structural evidences for possible integration of sensory and polymodal pathways onto PKCδ+ neurons, we asked what the possible downstream targets of these neurons are in the EAc. Both STLV and CeM, which are considered as the main outputs subnuclei of the EAc, have long been known as important intrinsic targets of STLD and CeL/C (Dong et al. 2001b; Cassell et al. 1999). It has been shown that PKCδ+ neurons in the CeL/C project to CeM (Haubensak et al. 2010; Li et al. 2013), but the neurochemical organization of connections inside the STL and between CeA and STL is still elusive. We thus injected the retrograde tracer CTb into the CeM (Fig. 6) or the STLV (Fig. 7), followed by triple fluorescent labeling for neuronal markers.

**Fig 6.**
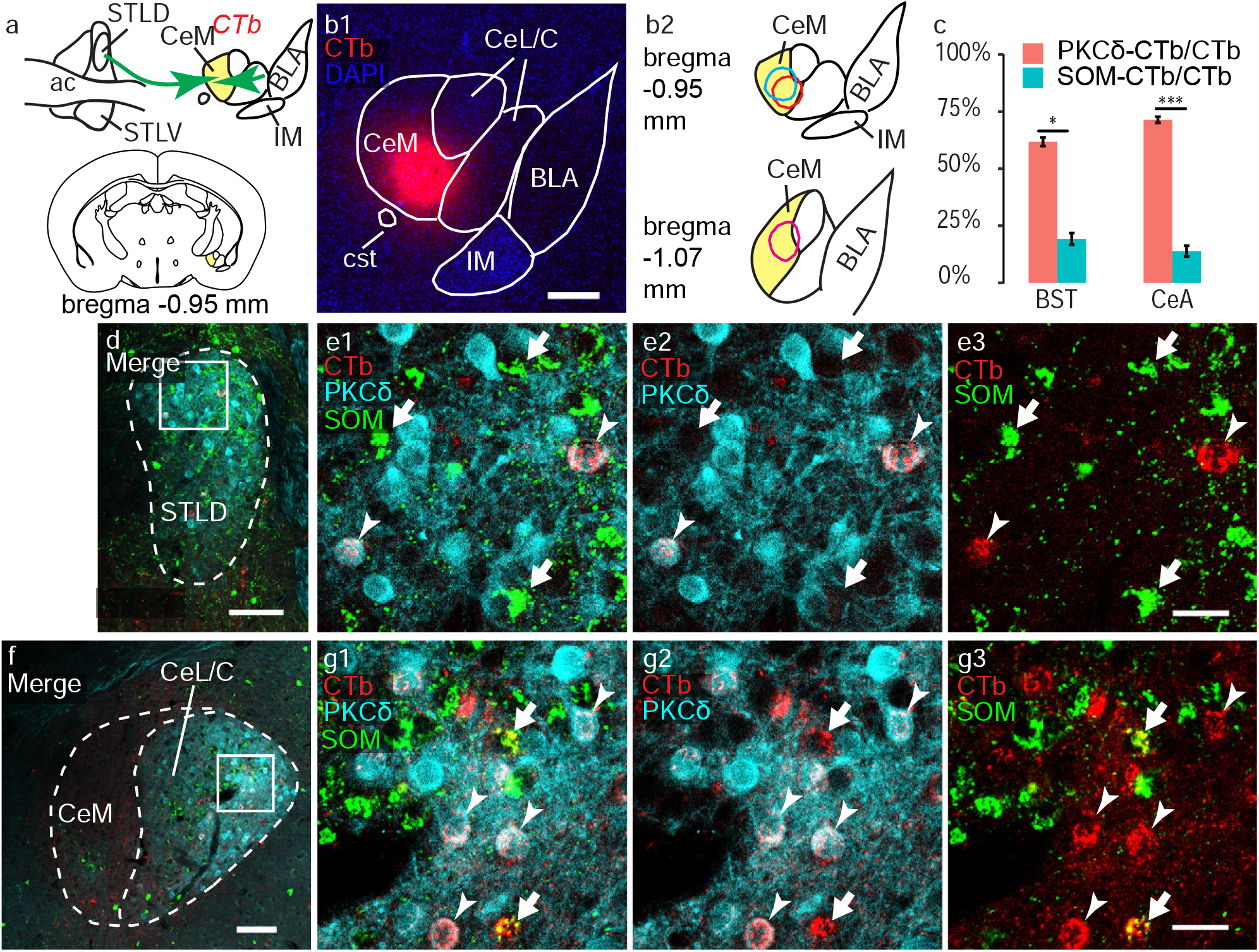
Retrogradely labeled CeM-projecting neurons in STLD and CeL/C express PKCδ. After injection of the retrograde tracer CTb into rostral CeM (a, b, bregma -0.95 mm), triple labeling of SOM (green), PKCδ (cyan) and CTb (red) was performed on STLD (d, e) and CeL/C sections (f, g). The injection sites (n = 3; b1, b2) were confined to the rostral CeM, with minimal extension into nearby regions. c Percentages of CTb+ somas positive for PKCδ and SOM in the STLD (PKCδ 60.8 ± 1.5 %; SOM 19.2 ± 2.6 %; two sample t-test, p < 0.05) and CeL/C (PKCδ 71.4 ± 1.3 %; SOM 13.9 ± 2.4 %; two sample t-test, p < 0.001). Confocal imaging in the STLD (d) and CeL/C (f) shows that retrogradely labeled CTb+ neurons were frequently PKCδ+ (e1, e2, g1, g2; arrowheads), but rarely SOM+ (e1, e3, g1, g3; short arrows). Abbreviations: see list. Scale bars: b1, 250 μm; d, 100 μm; e1 – e3, 25 μm; f, 100 μm; g1 - g3, 25 μm.

**Fig 7.**
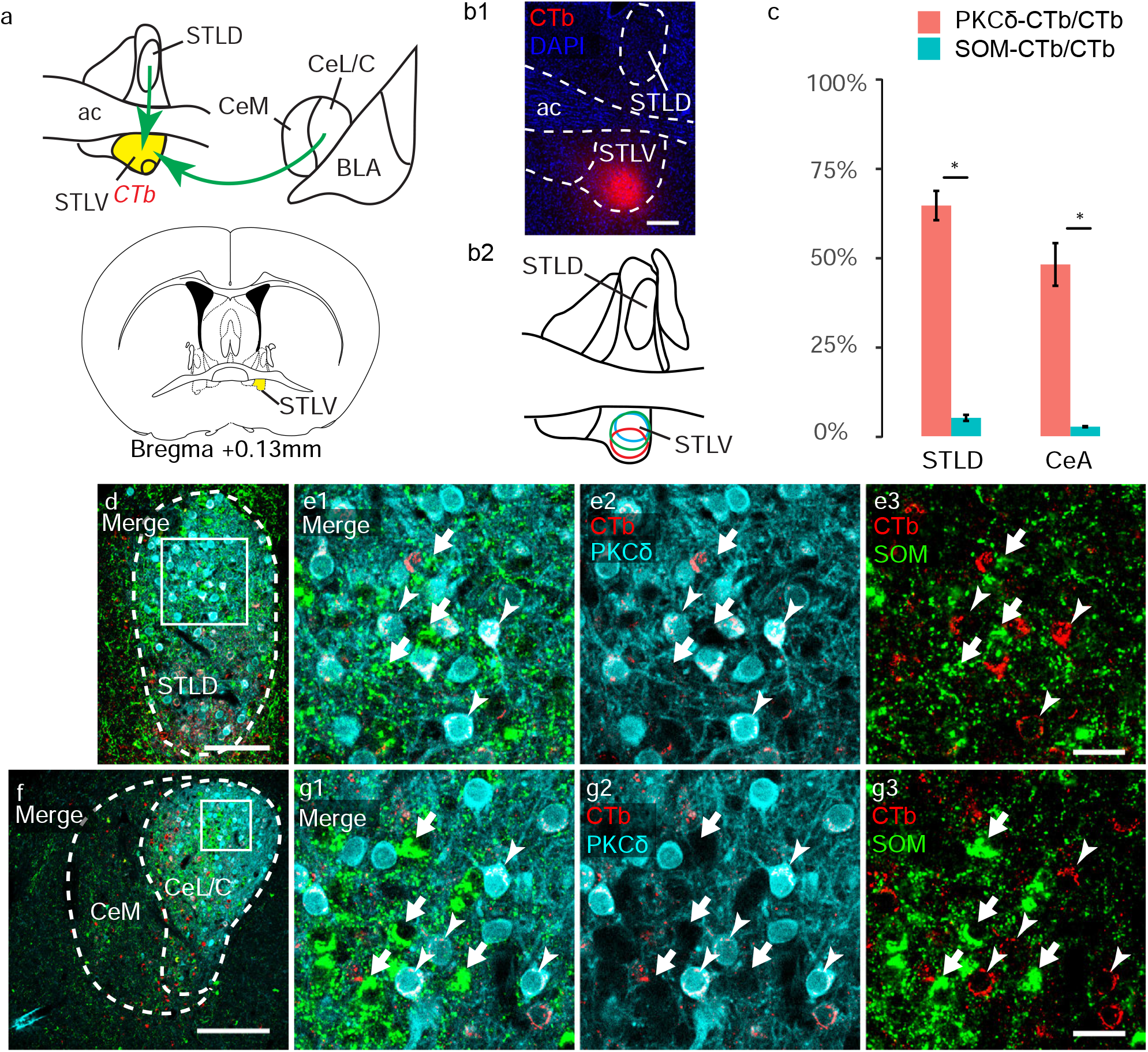
Retrogradely labeled STLV-projecting neurons in STLD and CeL/C express PKCδ. After injection of the retrograde tracer CTb into anterior STLV (a, b, bregma + 0.13 mm), triple labeling of SOM (green), PKCδ (cyan) and CTb (red) was performed on STLD (d, e) and CeL/C sections (f, g). The injection sites (n = 3; b1 – b2) were confined to the STLV with minimal extension to nearby areas. c Percentages of CTb+ somas positive for PKCδ and SOM in the STLD (PKCδ 64.6 ± 4.1 %; SOM 5.1 ± 0.1 %; two sample t-test, p < 0.05) and CeL/C (PKCδ 48.1±0.6 %; SOM 2.7 ± 0.2 %; two sample t-test, p < 0.05). Confocal imaging in the STLD (d) and CeL/C (f) shows that retrogradely labeled CTb+ neurons were frequently PKCδ+ (e1, e2, g1, g2; arrowheads), but rarely SOM+ (e1, e3, g1, g3; short arrows). Abbreviations: see list. Scale bars: b1, 200 μm; d, 100 μm; e1 – e3, 25 μm; f, 200 μm; g1 - g3, 25 μm.

CTb injections (n = 3) in rostral CeM (bregma level: -0.95/-1.07 mm) were centered in its ventral or dorsal portions (Fig. 6a), based on the cytoarchitectural features in DAPI staining (Fig. 6b) and the observation of typical retrograde labeling in rostral lateral amygdala (LA) and InsCx. In these cases, a robust retrograde labeling was found in the CeL/C (Fig. 6f, g), while much fewer cells were labeled in STLD (Fig. 6d, e). Quantitative analysis of the colocalization between CTb and PKCδ or SOM immunoreactivity revealed that, among the CeM-projecting neurons in CeL/C, 71.4 ± 1.3 % (Mean ± SEM) co-labeled with PKCδ and 13.9 ± 2.4 % with SOM (two sample *t*-test, p-value = 0.0009). In comparison, 60.8 ± 1.5 % of CTb+ cells in STLD were PKCδ+, but only 19.2 ± 2.6 % of them were SOM+ (two sample t-test, p-value = 0.002). CTb injections (n = 3) into STLV area (possibly including the fusiform nuclei) (Fig. 7a,b) revealed a considerable number of labeled neurons in STLD and CeL/C. The injection cores were confined to STLV as judged by DAPI staining and few/no retrograde labeling occurred in the STMA and medial amygdaloid nucleus (MeA). In STLD, we found that 64.6 ± 4.1% (Mean ± SEM) of CTb+ neurons were PKCδ+, while only 5.1 ± 0.1% of them were SOM+ (two sample *t*-test, p-value = 0.011). In CeL/C, 48.1±0.6% of STLV-projecting neurons were PKCδ+, by contrast only 2.7 ± 0.2 % were SOM+ (two sample *t*-test, p-value = 0.048). Taken together, our data suggest a significant role of PKCδ+ neurons in relaying information flow within EAc by connecting STLD and CeL/C with STLV and CeM. However, a sizeable part of the projections from STLD and CeL/C to STLV and CeM may originate in PKCδ-/SOM-neurons.

### Both PKCδ+ and SOM+ neurons are involved in STLD-CeL/C reciprocal connections

Although STL and CeA have been known to be reciprocally connected to each other (Dong et al. 2001a; Gungor et al. 2015; Sun et al. 1994; Sun and Cassell 1993; Sun et al. 1991), it remains not clear which cell types mediate the mutual connections between STLD and CeL/C. In mouse, rabies virus tracing from CeL PKCδ+ neurons revealed a dense neuronal labeling in dorsal STL (Cai et al. 2014), which arose an interesting speculation that PKCδ+ cells might serve as intrinsic projection neurons between STLD and CeL/C. To test this hypothesis, we carried out retrograde (Fig. 8) and anterograde (Fig. 9) tracings from STLD and CeL/C, followed by immunostaining of the tracers, PKCδ and SOM.

**Fig 8.**
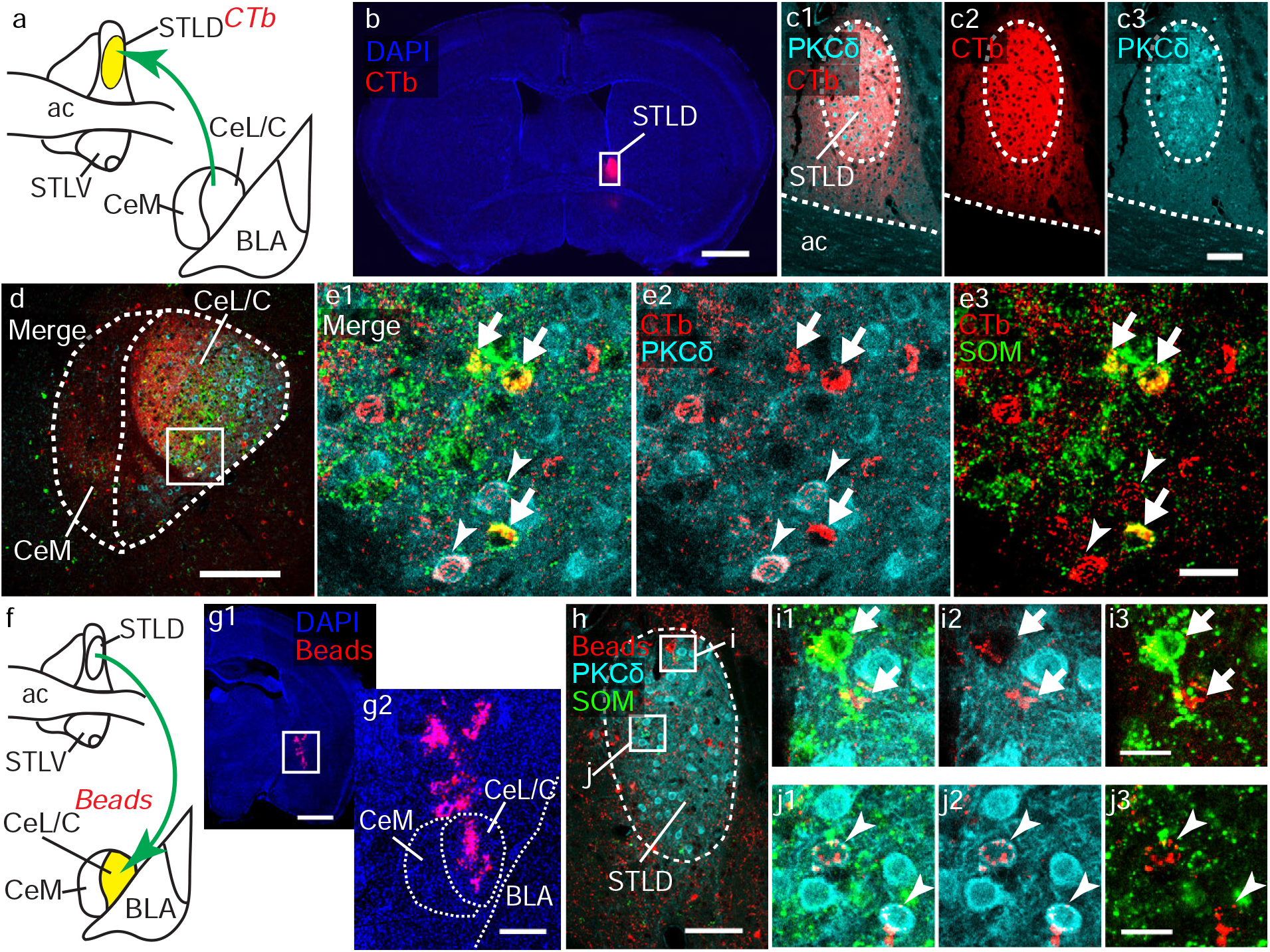
Retrogradely labeled STLD- and CeL/C-projecting neurons express PKCδ or SOM. Following by CTb injection in STLD (a – c, bregma level + 0.13 mm) or red retrobeads in CeL/C (f, g, bregma level – 1.43 mm), triple immunofluorescence labeling was carried out for CTb (red), PKCδ (cyan) and SOM (green), while intrinsic fluorescence from retrobeads was used. In STL, CTb injection site was limited to the PKCδ-expressing STLD (b, c). In ipsilateral caudal CeL/C (c), confocal image (z stack = 5.78 μm) identified CTb+/PKCδ+ colabeled neurons (arrowheads) and CTb+/SOM+ ones (short arrows) (el – e3). Pressure injection of red retrobeads resulted in dense deposit in CeL/C (gl, g2). Subsequent colocalization analysis revealed double labeling from SOM+ populations (arrowheads) (il - i3) and PKCδ+ ones (short arrows) (jl – j3). Abbreviations, see list. Scale bars: b, 1000 μm; cl - c3, 100 μm; d, 200 μm; el – e3, 25 μm; gl, 1000 μm; g2, 250 μm; h, 100 μm; il – i3, 20 μm; jl – j3, 20 μm.

**Fig 9.**
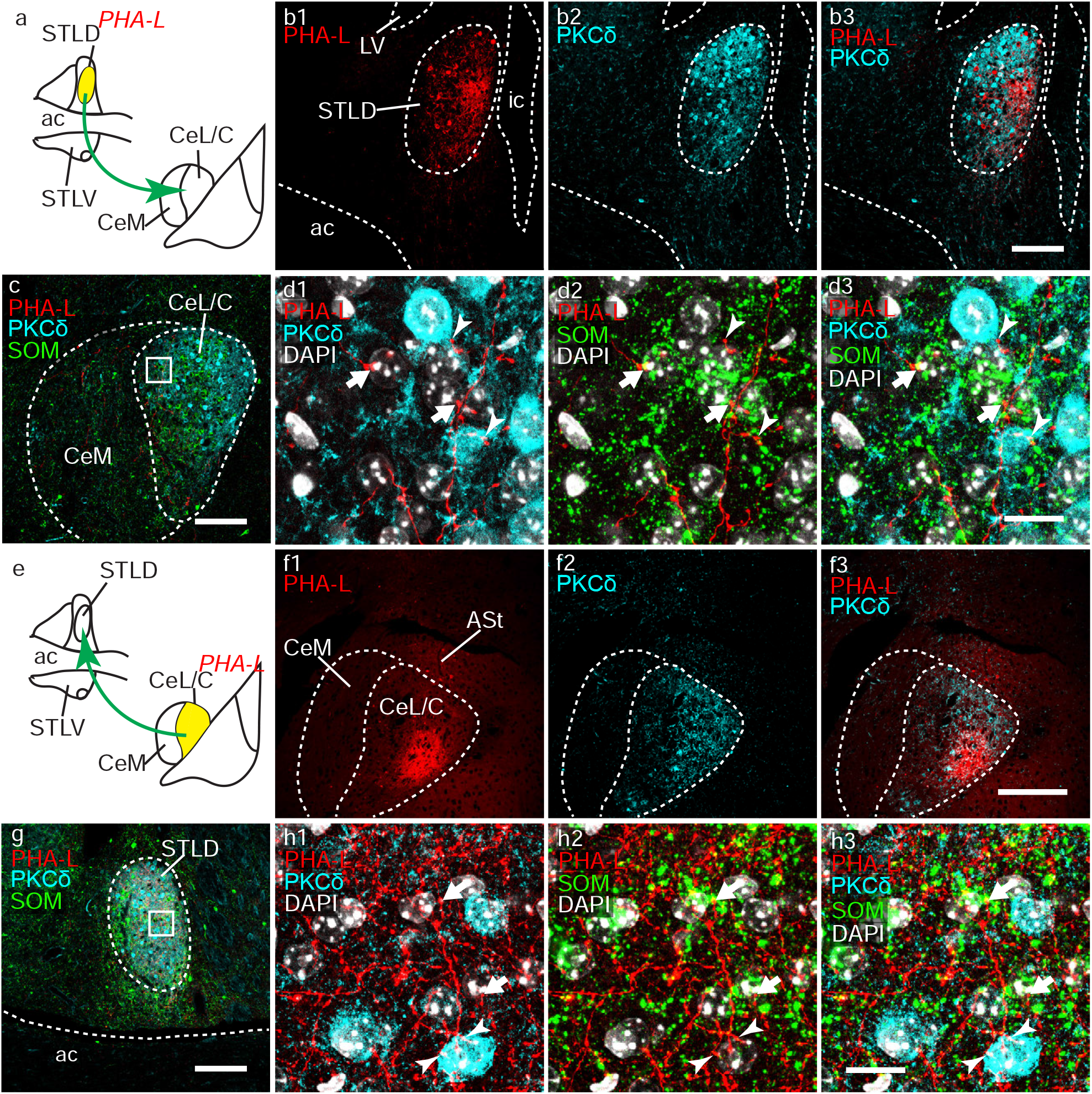
Anterogradely labeled STLD or CeL/C axonal projections can target both PKCδ+ and SOM+ neurons. Following by PHA-L injection in STLD (a, b; bregma level + 0.01 mm) and in CeL/C (e, f; bregma level – 1.55 mm), triple immunofluorescence labeling was carried out for PHA-L (red), PKCδ (cyan) and SOM (green). In STL, restricted PHA-L injection site was confined to the STLD (b1 – b3). In caudal level of CeL/C (c), confocal imaging (z stack = 11.9 μm) revealed PHA-L+ varicosities apposed to PKCδ+ (arrowheads) and SOM+ neurons (short arrows) (dl – d3). In another case, PHA-L injection into CeL/C (fl – f3) resulted in dense axonal labeling in STL, especially in STLD (g). With high magnification confocal images (z stack = 10.9 μm), PHA-L+ varicosities were observed forming close apposition with PKCδ+ (arrowheads) and SOM+ (short arrows) (hl – h3). Abbreviations, see list. Scale bars: b, 150 μm; c, 150 μm; d1 – d3, 15 μm; f1 – f3, 200 μm; g, 150 μm; h1 – h3, 15 μm.

To determine if PKCδ+ and/or SOM+ neurons in CeL/C project to STLD, CTb injections were done in the STLD (n = 2; bregma level +0.13 mm). The injection sites were restricted to the PKCδ-expressing STLD (Fig. 8 b – c) and led to a large number of retrogradely labeled neurons in CeM and CeL/C, while labeling in medial amygdala was rarely seen (Fig. 8d). With confocal analysis, we found both CTb+/PKCδ+ and CTb+/SOM+ double labeled neurons in ipsilateral CeL/C (Fig. 8e). In a similar attempt, we labeled CeA-projecting neurons in STLD by injecting retrobeads into caudal CeL/C (n = 2; Fig. 8f, g). Here, the retrobeads were preferred to CTb to avoid any leakage in the CeM. Retrobeads indeed produced a local injection zone in CeL/C, without extension into CeM (Fig. 8g). Despite a leakage along the micropipette track into the amygdalostriatal transition area (ASt) and globus pallidus (GP), we considered that possible confounding retrograde labeling in STLD would be negligible as anterograde tracing from STLD rarely labeled axons in ASt region. In this case, similar to that of CeL/C, the retrograde labeling could be found in both PKCδ+ neurons (Fig. 8j) and SOM+ neurons (Fig. 8i).

Thus, our evidences indicate that both PKCδ+ and SOM+ neurons contribute to intra-EAc connections, mediating mutual talks between the STLD and CeL/C. To further identify the possible neurochemical profile of the neurons that receive inputs from STLD or CeL/C, we injected PHA-L in STLD or CeL/C and looked for potential appositions of anterogradely labeled axons with PKCδ+ and SOM+ neurons (Fig. 9).

Small PHA-L injections into STL (n = 1) produced a restricted labeling of neurons and processes, confined to the PKCδ-expressing STLD (Fig. 9b). In caudal CeA, a moderate density of PHA-L+ axonal branches and terminals were found in CeM and CeL/C (Fig. 9c). Confocal images (z stack = 11.9 μm) at high magnification showed that PHA-L+ varicosities from single continuous axons ramifications could be found apposed to both PKCδ+ and SOM+ neurons (Fig. 9d). Similarly, PHA-L injection sites into caudal CeL/C were centered in CeL/C, without leakage in BLA or CeM (n = 1; Fig. 9f). Numerous PHA-L+ axons could be observed in STL, with the highest density in the STLD (Fig. 9 g). Apposition analysis following triple immunofluorescence staining revealed that many axon terminals formed close appositions with PKCδ+ and SOM+ neurons (Fig. 9h).

Thus, we concluded that projections from PKCδ+ and SOM+ neurons in STLD and CeL/C can target both PKCδ+ and SOM+ in the same subdivisions.

### SOM+ neurons in STLD and CeL/C are the main sources of downstream projections to brainstem

Apart from the intra-EAc projection, neurons in STLD and CeL/C give rise to efferent to extra-EAc targets as well, including, the LPB and the PAG (Tokita et al. 2009; Dong et al. 2001b; Petrovich and Swanson 1997; Gray and Magnuson 1992; Moga and Gray 1985). Interestingly, brainstem-projecting neurons in STL and CeA share similar neuropeptidergic features in rats (Moga et al. 1989). In mice, it has been shown that SOM+ cells in CeL/C project to PAG (Penzo et al. 2014). In order to establish the neurochemical identity of neurons in STLD and CeL/C projecting to brainstem, we injected retrograde tracers into LPB and PAG.

Fluorogold (FG) injections in LPB (n = 3; Fig. 10) usually resulted in minor lesion centered within LPBE (bregma -5.19 mm) and diffuse expansion into other subdivisions of LPB (Fig. 10b). The retrograde labeling in ST and amygdala was specifically restricted to STL and CeA, especially in STLD and CeL/C, with much sparser labeling in STLP and CeM. In STLD (Fig. 10d, e) as in CeL/C (Fig. 10f, g), numerous FG+ cells were SOM+ but very few were PKCδ+. Quantitative analysis (Fig. 10c) revealed that, SOM+ neurons accounted for 62.7± 0.4 % and 63.9± 0.7 % of the retrogradely labeled cells in STLD and CeL/C, respectively, whereas only 6.1± 0.4 % and 6.9±0.7 %; of FG+ neurons were PKCδ+ (two sample t-test, STLD p-value = 0.011, CeL/C p-value = 5.37e-06).

**Fig 10.**
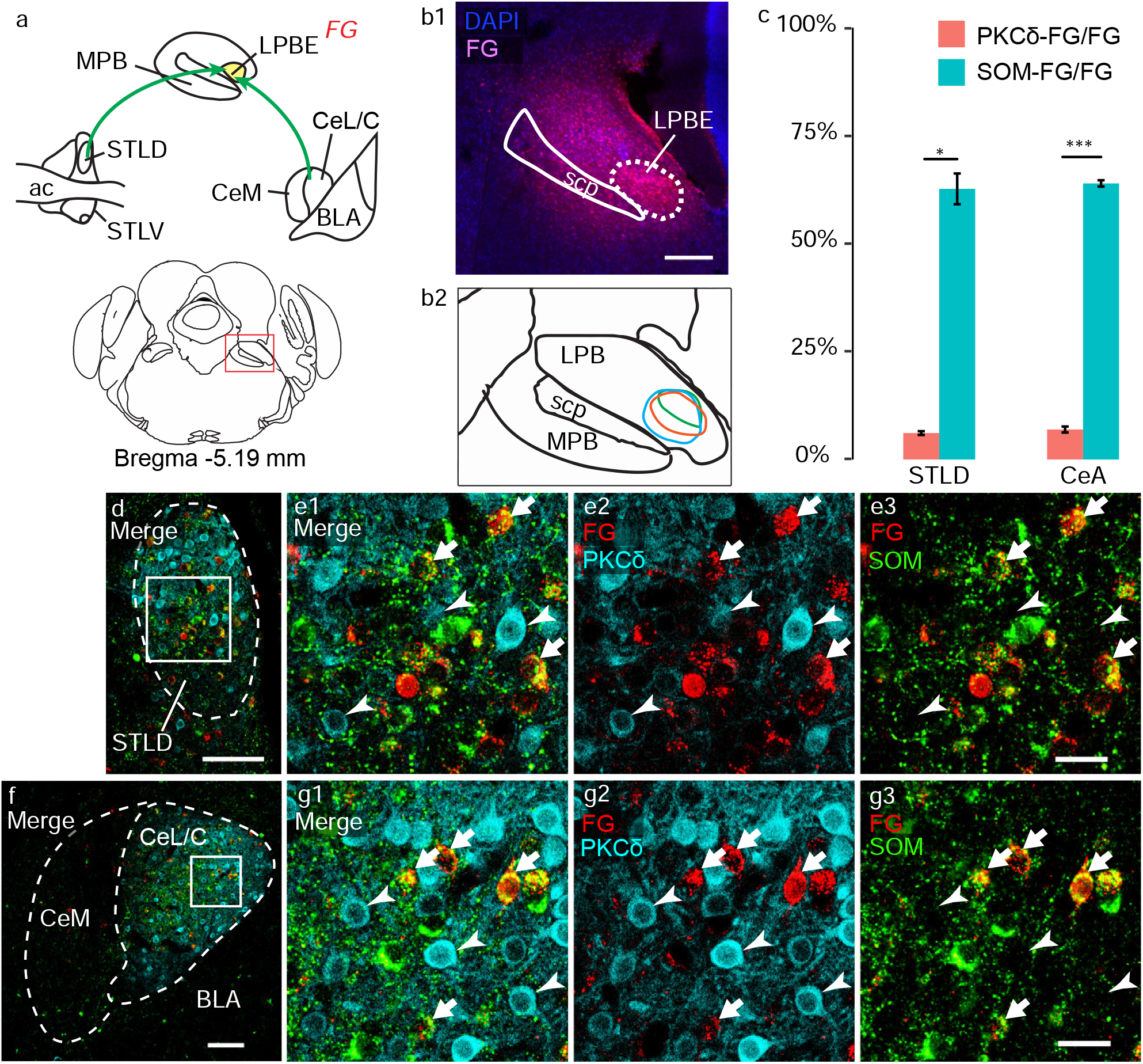
LPBE-projecting neurons in STLD and CeL/C express mainly SOM. Triple labeling of FG (red), PKCδ (cyan) and SOM (green) in STLD (d, e) and CeA (f, g) was performed after FG retrograde tracing from LPBE (a, b, bregma level – 5.19 mm). The FG injection sites (n = 3) were centered in LPBE, with minor diffusion in other LPB subdivisions (b1, b2). c Percentage of FG+ somas positive for PKCδ and SOM in STLD (PKCδ, 6.1± 0.4 %; SOM, 62.7± 0.4 %; p-value < 0.05) and in CeL/C (PKCδ, 6.9±0.7 %; SOM, 63.9± 0.7 %; p-value < 0.001). d - g Confocal images show rare colabeling of PKCδ (arrowheads) with FG, whereas SOM+ neurons (short arrows) frequently contained FG, in both STLD (d, e) and CeL/C (f, g). Abbreviations, see list. Scale bars: b1, 250 μm; d, 100 μm; e1 - e3, 25 μm; f, 100 μm; g1 - g3, 25 μm.

To further examine the possibility that STLD and CeL/C projections to LPBE can target CGRP+ neurons, we processed sections from animals with PHA-L injections into STLD (same case as in Fig. 9b) or into CeL/C (same case as in Fig. 9f), to label PHA-L and CGRP on LPB sections Consistent with the previous retrograde tracing, intense labeling of PHA-L+ axons was observed in LPB, especially dense in LPBE, following PHA-L injection in STLD (Fig. 11b) or CeL/C (Fig. 11d). CGRP+ neurons were concentrated in the ventrolateral part of the LPB, including the LPBE. Confocal analysis at high magnification revealed frequent, although not exclusive, appositions between PHA-L+ axonal varicosities, from STLD and CeL/C, and LBPE somas containing CGRP immunofluorescence. (Fig. 11b, d).

**Fig 11.**
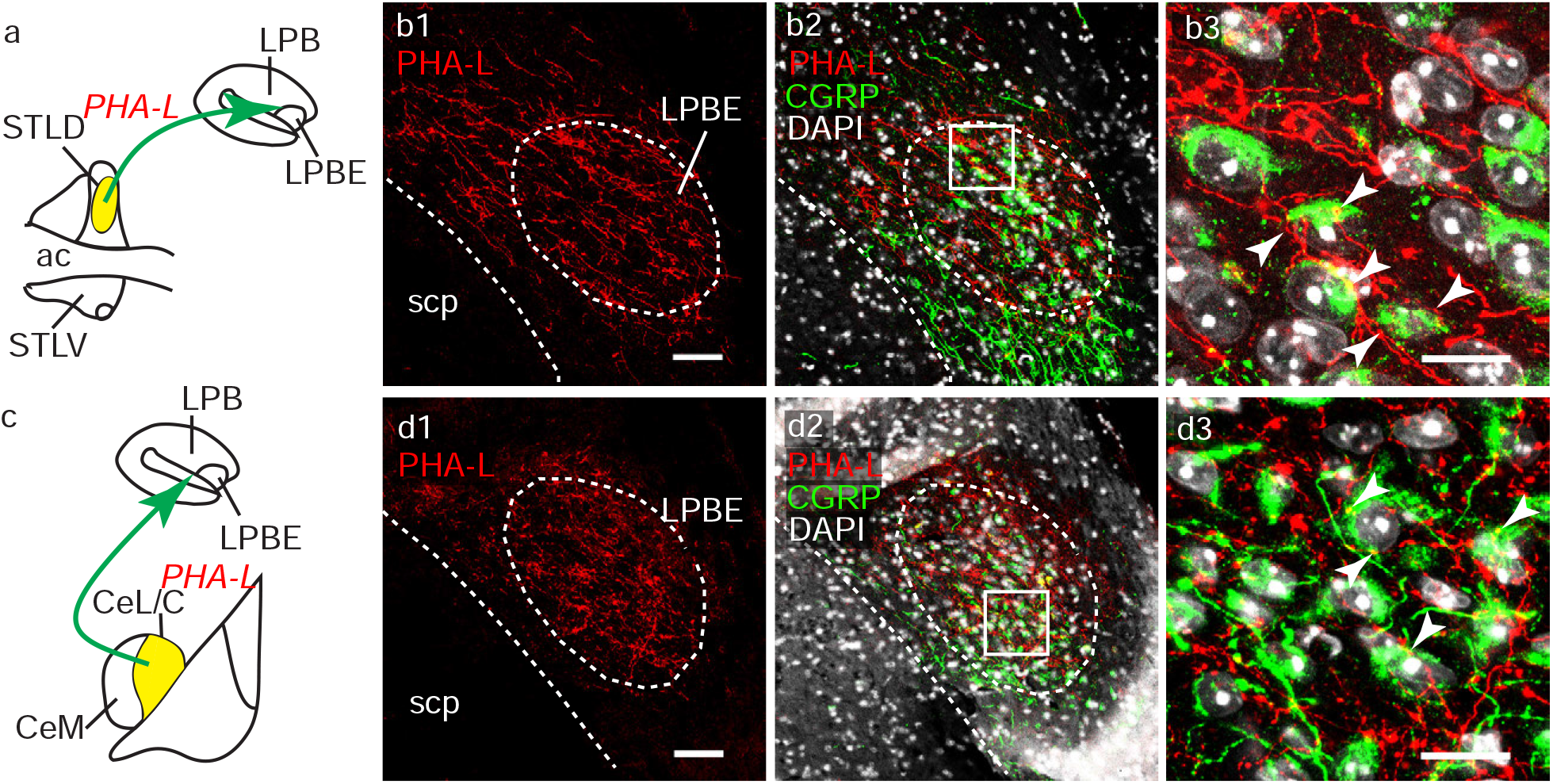
STLD and CeL/C projections can target CGRP+ neurons in LPBE. Double immunofluorescent labeling for PHA-L (red) and CGRP (green), together with DAPI (white) in LPB, after PHA-L injection in STLD (a) or CeL/C (c). A dense PHA-L+ axonal labeling was observed in LPB, especially LPBE where it overlapped with the presence of CGRP+ neurons (b1, b2, d1, d2). With high magnification confocal images, axonal apposition onto CGRP+ soma (arrowheads) were frequently observed for projections from STLD (b3; z stack = 9 μm) and CeL/C (d3; z stack = 7.9 μm). Abbreviations, see list. Scale bars: b1 – b2, 50 μm; b3, 15 μm; d1 – d2, 50 μm; d3, 15 μm.

To investigate the EAc projection to PAG, we used retrograde tracers FG or CTb and performed triple immunofluorescence staining for the tracer, PKCδ and SOM. In order to achieve reasonable number of retrograde labeling in STLD and CeL/C (versus STLP or CeM), we produced large injection sites with tracer deposits extending into the lateral (LPAG) and ventrolateral (VLPAG) columns of the PAG and dorsal raphe nuclei (DR; bregma -4.47/-4.59 mm; Fig. 12b, c). Retrogradely labeled cells were found in STL and CeA, including STLD (Fig. 12d) and CeL/C (Fig. 12f). While no quantification has been done (one FG case and one CTb case), we observed that more than half of the retrogradely labeled neurons colocalized with SOM immunofluorescence in STLD (Fig. 12e) and in CeL/C (Fig. 12g), but almost never with PKCδ signal. These data indicate, in both STLD and CeL/C, SOM+, but not PKCδ+ neurons, project to PAG/DR areas.

**Fig 12.**
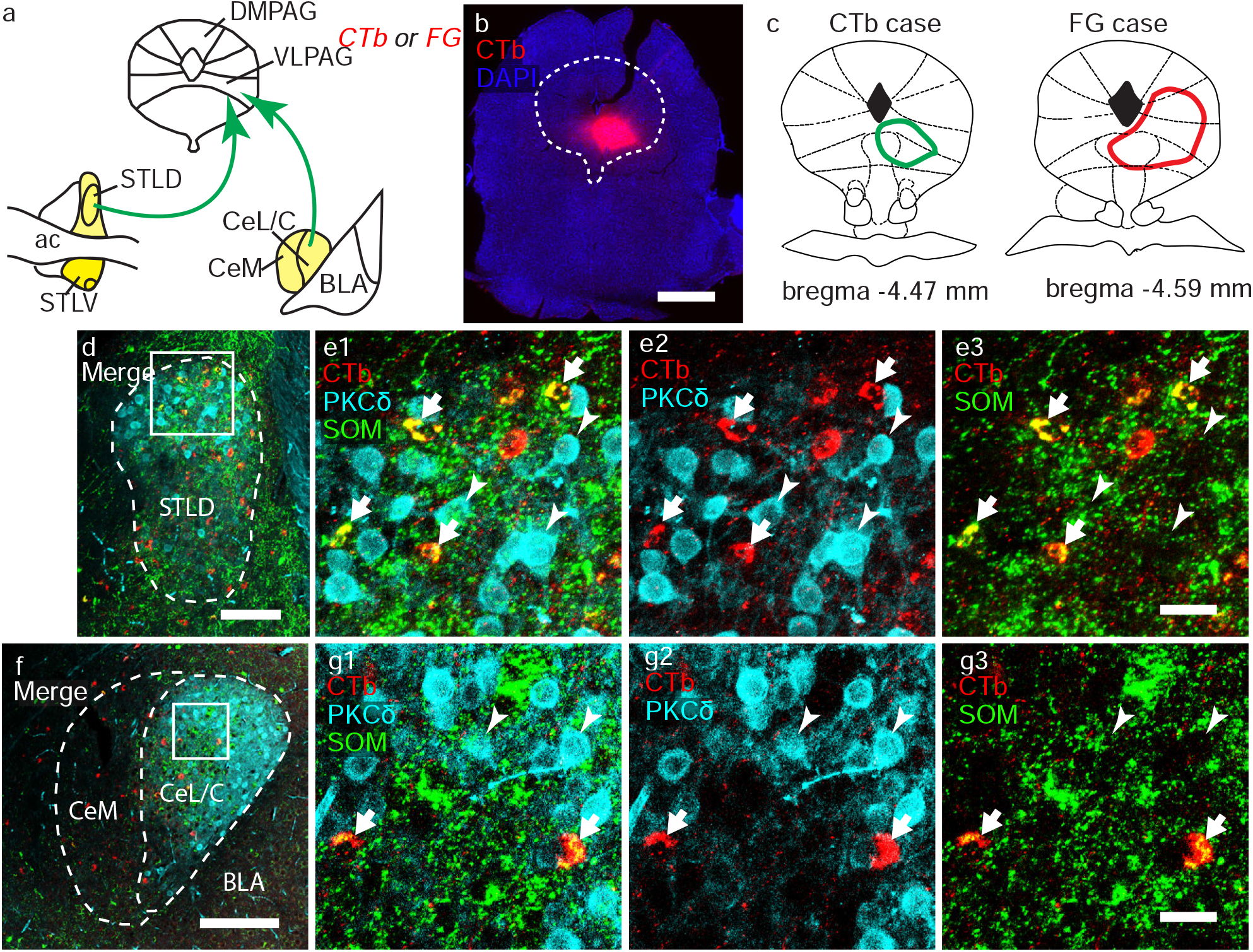
PAG/DRN-projecting neurons in STLD and CeL/C express SOM. Triple labeling of CTb (red), PKCδ (cyan), and SOM (green) in STLD (d, e) and CeA (f, g), after CTb injection into PAG areas (a-c). b illustrates a CTb injection site. c The CTb and FG injection sites covered lateral (LPAG), ventrolateral PAG (VLPAG) and dorsal raphe (DR). d, g Confocal images show that most of the CTb+ neurons were colabeled by SOM (short arrows) in STLD (e1, e3; z-stack = 15.8 μm) and CeL/C (g1, g3; z-stack = 5.9 μm), but not PKCδ (arrowheads) in both areas (e1, e2, g1, g2). Abbreviations, see list. Scale bars: b, 1000 μm; d, 100 μm; e1 – e3, 25 μm; f, 200 μm; g1 – g3, 25 μm.

Taken together, these data supports a major role of STLD and CeL/C SOM+ neurons in mediating long range projections to LPB and PAG, while PKCδ+ neurons contribute very little to this outputs.

## DISCUSSION

In this study, we addressed the possibility of a similar organization of specific cell-type neuronal circuits in STLD and CeA of mice, by combining retrograde and anterograde tract-tracing with immunofluorescent staining. Overall, we looked at three different aspects of neuronal circuit organizations of EAc, including the long-range inputs, intrinsic projections and long-range external outputs. We propose a model of cell-type specific parallel microcircuits in EAc, based on the connectivity of PKCδ+ and SOM+ neuronal populations (Fig. 13).

**Fig 13.**
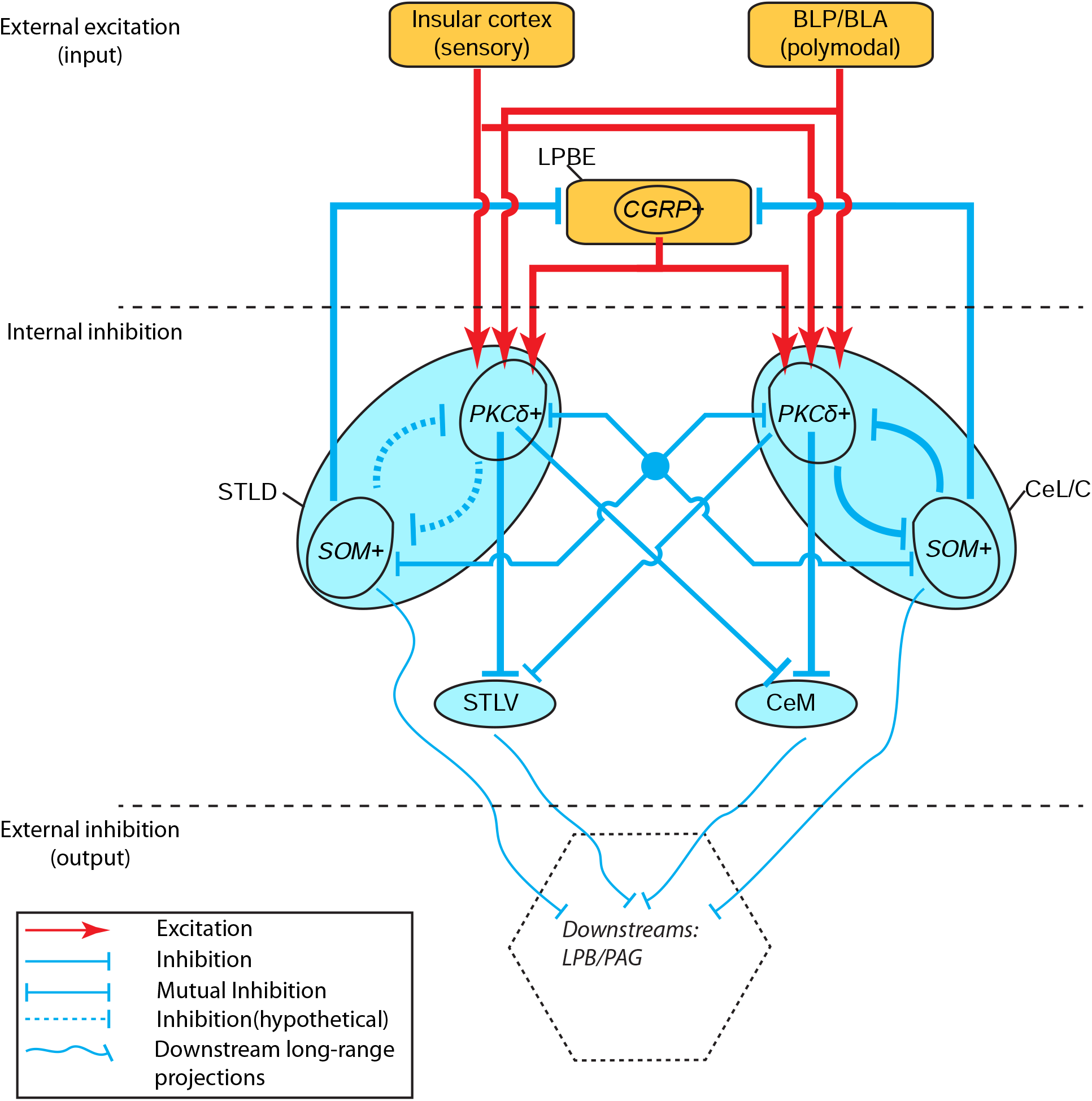
A simplified model of parallel, cell-type specific, neuronal circuits in EAc. This model highlight the similar configuration of cell-type specific neuronal circuits in STL and CeA, based on PKCδ+ neurons and SOM+ neurons in STLD and CeL/C. Excitations coming from InsCx or basolateral amygdala nuclei, together with CGRP inputs from LPB, can converge onto PKCδ+ neurons in STLD and CeL/C, in a similar fashion. The internal inhibitory circuits in STLD or CeL/C are probably mediated by PKCδ+ and SOM+ neurons.. The internal long-range projections to STLV and CeM are primarily mediated by PKCδ+ neurons, while mutual connection between STLD and CeL/C can be mediated by both types, although the connection from CeL/C to STLD is stronger than the in reverse direction. The external inhibition to LPB and PAG can be mediated by SOM+ neurons in STLD or CeL/C, as well as undefined populations in STLV and CeM.

For the external inputs, our data support the hypothesis that multiple excitatory inputs can converge onto single neuronal populations in STLD and CeL/C. For instance, excitatory sensory information from cortex or polymodal information from amygdala nuclei (i.e. BLP) can converge to PKCδ+ neurons which at the same time are innervated by excitatory CGRP+ sensory input from brainstem (i.e. LPB).

These excitatory drives onto distinct neuronal populations in EAc are then processed by intrinsic inhibitory circuits, including local (i.e. SOM+ ➜SOM+ in CeL/C) (Hunt et al. 2017; Douglass et al. 2017) and long-range connections (i.e. PKCδ+ neurons in CeL/C ➜STLV). Because much less is known on local inhibitory circuits in STLD, we hypothesize that a similar configuration also exists there (dashed line, Fig. 13), featured with both homotypic (i.e. SOM+ ➜ SOM+, not shown) and heterotypic (SOM+ ➜ PKCδ+) connections (Fig. 13). For intrinsic long-range connections, we confirmed similar preferential innervations of STLV and CeM by PKCδ+ neurons in STLD and CeL/C, although sparse innervations from SOM+ populations are observed. The long-range, mutual, connections between STLD and CeL/C can be carried out by both types of neurons.

Finally, information from EAc are carried out mainly by SOM+ neurons in STLD and CeL/C, as well as by undefined neuronal groups in STLV and CeM. Notably, we find that SOM+, not PKCδ+, populations, mediate the output to LPBE and PAG areas, and possibly to other known downstream targets.

### Technical considerations

In this study, the quality of injection sites are critical for reliable and accountable explanations drawn from tract-tracing experiments. In total, we used FG/CTb and retrobeads for retrograde tracing, PHA-L/BDA for anterograde tracing. After checking the neuroanatomical localization of injection sites on successive coronal brain sections, we excluded cases with confounding spillovers. When applied by iontophoresis, CTb, BDA and PHA-L reliably produced limited injection sites, usually confined to the limits of the target nucleus (i.e. see CTb injection into STLD, Fig. 8). Iontophoresis of FG into LPBE usually resulted in strong diffusive labeling in the other subdivisions of LPB, but we found minimal contaminations from these non-LPBE subdivision as suggested by minimal retrograde labeling in non-EAc subdivisions. In our hands, pressure injection of retrobeads in CeL/C usually resulted in well defined injection sites, but also in tracer deposits along the pipette track, in ASt, GP or CPu. However, none of these areas are innervated by STLD, based on the litterature (Weller and Smith 1982; McDonald 1991) and on our results from retrograde tracing.

We relied on antibodies to determine the cellular identity of PKCδ+ and SOM+ neurons. Due to unknown reasons, we observed that the immunofluorescent signal in STLD and CeL/C was weaker than the one in thalamic PKCδ+ neurons and cortical or striatal SOM+ neurons in the same brain sections. Nevertheless, the primary antibody for PKCδ we used was shown to detected most of the PKCδ-cre-positive neurons in a transgenic mouse line (Haubensak et al. 2010). The antibody against SOM gave a specific labeling of SOM-expressing neurons (Jhou et al. 2009) but seems to reveal much less neurons than what is observed in SOM-cre mouse line (Li et al. 2013). Finally, CGRP antibody revealed terminal fields in EAc that are largely consistent with previous reports (Dobolyi et al. 2005). Thus, these tools allowed showing the basket-like CGRP+ axon terminals in appositions with PKCδ+ neurons in STLD and CeL/C, as PKCδ+ signal usually reliably labeled the whole cell body and proximal dendrites. We very rarely observed these obvious basket-like terminals around SOM-expressing neurons. However, we can not exclude that more discreet CGRP+ terminals might contact SOM+ neurons at the level of soma or dendrites. For the same reasons, we also likely underestimated the extent of CGRP+ contacts with PKCδ+ neurons as non-basket CGRP+ varicosities apposition with neurons is difficult to ascertain in our experimental conditions. The confirmation of CGRP+ synaptic contact by immunostaining of presynaptic markers or by synaptic ultra-structures with electronic microscopy would be necessary.

### Neurochemical features of EAc

The subdivisions of EAc have long been known to express a variety of neuropeptides and receptors, such as ENK, CRF, SOM, dopamine receptors, serotonin receptors 2a (Htr2a) (Cassell et al. 1986; Cassell et al. 1999; De Bundel et al. 2016; Douglass et al. 2017; Veinante et al. 1997). In this study, we focused on the cellular connectivity of PKCδ+ and SOM+ neurons, primarily because these two neuronal populations are largely non-overlapping and constitute the majority of local GABAergic neurons in CeA (Haubensak et al. 2010; Li et al. 2013). Using double immunofluorescent staining, we found similar segregation and expression patterns of PKCδ and SOM in CeA than in previous reports on cre mouse line (Li et al. 2013). In addition, we describe for the first time a similar pattern in STLD. Eventhough immunofluorescent staining associated to the highly sensitive CARD method (Hunyady et al. 1996) allows us to visualize many SOM+ and PKCδ+ neurons in EAc, the transgenic mouse lines might provide a more robust and reliable way to label these neurons (Li et al. 2013).

On the other hand, these two neuronal populations can intersect with other neuronal markers. For example, More than 70% of PKCδ neurons in CeL/C and STLD coexpressed the dopamine receptor D2 (D2R) in a Drd2-cre-EGFP mouse (De Bundel et al. 2016). PKCδ neurons do not overlap with Htr2a-expressing cells in CeL, but more than half of Htr2a+ neurons coexpress SOM (Douglass et al. 2017). SOM+ neurons in both STL and CeA can also coexpress neuropeptide Y (NPY) (Wood et al. 2016). Thus, it is possible that some of the EAc PKCδ+ or SOM+ neurons revealed in this study also belong to other specific neuronal populations.

### Comparison with other studies on cell-type specific circuits in EAc

#### Long-range inputs

The identity of presynaptic inputs from extra-EAc sources have been studied in various ways and are in accordance with our study. Projection neurons from LPBE and BLP are the best studied compared to insular cortex.

CGRP+ neurons in LPBE have been shown project to CeL/C or STL by immunohistochemistry (Dobolyi et al. 2005), retrograde tract-tracing (Carter et al. 2013), cell-type specific rabies tracing (Cai et al. 2014) and optogenetic mapping (Carter et al. 2013; Sato et al. 2015). Furthermore, CGRP receptor (CGRPR)-expressing CeL/C neurons were proved to be innervated by LPBE CGRP+ neurons, using a double cre mouse line (Han et al. 2015), while the connectivity of CGRPR+ neurons in STL remains relatively unexplored. In our study, most of the CGRP+ terminals in CeL/C and STLD, as well as many of the axon terminals anterogradely labeled from LPBE, appear as basket perisomatic terminals, morphologically similar to those described in studies on rat (Sarhan et al. 2005; Dobolyi et al. 2005) and mouse (Campos et al. 2016). We found a preferential targeting of CGRP+ terminals onto PKCδ+ soma and proximal dendrites, but not SOM+ ones. However, we cannot exclude a synaptic or extra-synaptic influence of CGRP on SOM+ neurons, as a recent study indicates that only about half of calcitonin receptor-like (Calcrl) positive neurons coexpress PKCδ in CeC of mice (Kim et al. 2017).

The projection from the basolateral nucleus of the amygdala (BL) to CeA and STL, has been revealed by anterograde tract-tracing (Pitkanen et al. 1995; Dong et al. 2001a; Savander et al. 1996), monosynaptic rabies virus tracing (Kim et al. 2017) and optogenetic mapping (Li et al. 2013). It is worth noting that these CeA-projecting neurons are distributed differently along the rostral-caudal axis of the BL. Most of the CeA-projecting neurons are located in the caudal BL and express the protein phosphatase 1 regulatory subunit 1B (Ppp1r1b+); while a minority, projecting essentially to CeC expresses R-spondin 2 positive (Rspo2+) (Kim et al. 2017). In line with these findings, we found that CTb tracing from STLD and CeL/C resulted in dramatically more labeling in BLP than BLA. Insular cortex inputs to CeA and STL have also been previously described (Yasui et al. 1991a; McDonald et al. 1999; Sun et al. 1994) and have been shown to arise mainly from agranular and dysgranular areas. In this study, we provide further evidences supporting a convergence of long-range pathways onto individual PKCδ+ neuron in both STLD and CeL/C. However, this connection is not exclusive as we also observed axon terminals from BLP or insular cortex apposed to PKCδ-soma, which could be also targeted by non-CGRP LPBE projections. It also important to note that, if we show that a number of given inputs can converge onto PKCδ+ population, other inputs might favor different populations. For example, afferents from the thalamic paraventricular nucleus target two times more the SOM+ neurons than PKCδ+ neurons in CeL/C (Penzo et al. 2015).

#### Intrinsic circuits

In this study, information concerning the local intrinsic connections (i.e. inside STLD or CeL/C) are obscured by bulk tract-tracing method. However, recent works using genetic tools revealed complex disinhibitory circuit between PKCδ+ and SOM+ neurons in the CeA. (Ciocchi et al. 2010; Haubensak et al. 2010; Li et al. 2013; Janak and Tye 2015; Hunt et al. 2017; Douglass et al. 2017). Into the CeL/C, it has been shown that PKCδ+ neurons project to PKCδ-negative ones (Haubensak et al. 2010; Douglass et al. 2017) and non-PKCδ neurons project more to non-PKCδ cells (Hunt et al. 2017). By taking advantage of rabies virus tracing in multiple cre mouse lines, Kim and colleagues revealed a surprising complexity in the connections between several neuronal populations in CeL, including PKCδ, SOM, CRF, neurotensin, and tachykinin 2 (Kim et al. 2017). Here again, information on STLD local circuits is still missing.

On the other hand, connectivity between EAc subdivisions, including short-range ones linking CeA or STL subdivisions and long-range ones between CeA and STL subdivisions, can be well-resolved by restricted injection of retrograde tracer like CTb. Li and colleagues reported that about only 15% of CeM-projecting neurons are SOM+ in CeL/C (Li et al. 2013), while the majority of PKCδ+ neurons projects to CeM (Ciocchi et al. 2010; Li et al. 2013; Oh et al. 2014), which is consistent with the present findings. In comparison, limited information is available on STLV-projecting CeL/C or STLD neurons. We found that, similarly to the CeL/C-CeM pathway, PKCδ+ neurons are the main source of projection from the STLD to STLV with only a small contribution of SOM+ neurons. In addition we also evidenced the fact that similar proportion of these neuronal populations contribute to long-range projections from CeL/C to STLV and from STLD to CeM. It would be interesting to verify whether a single CeL/C or STLD neuron can project to both CeM and STLV, as it has been suggested for CeL neurons in rats (Veinante and Freund-Mercier 2003).While PKCδ+ neurons are clearly involved in these intrinsic EAc connections, it is worth noting that, in our hands, only 80% of CeM-projecting neurons in STLD or CeL/C can be attributed to PKCδ and SOM population, while about 30 – 50% of STLV-projecting ones were not labeled by either of the two markers. This suggests that other neuronal populations can significantly contribute to the EAc intrinsic long-range projection, especially to STLV. Indeed, other neurochemically defined neuronal populations have been shown to mediate mutual or unidirectional connection between STL and CeA, such as NPY+ (Wood et al. 2016), Htr2a+ (Douglass et al. 2017) and CRF+ populations (Pomrenze et al. 2015).

The cellular identity of the neurons mediating STLD – CeL/C mutual connections is also elusive. We showed by retrograde and anterograde tracing that both PKCδ+ and SOM+ neurons can be the sources and the targets of the STLD-CeL/C connections (Fig. 9). We also observed that retrograde labeling in STLD was much weaker than that in CeL/C, indicating a preferential CeL/C ➜ STLD direction..

#### Long-range outputs

EAc neurons projecting to LPB has been suggested to contain several different neuronal markers such as CRF, neurotensin, ENK and SOM (Moga et al. 1989; Panguluri et al. 2009; Magableh and Lundy 2014; Moga and Gray 1985). While PKCδ+ neurons have been demonstrated by optogenetic mapping to not, or faintly, project to LPBE (Douglass et al. 2017; Oh et al. 2014), a strong terminal field from CeL/C PKCδ+ neurons was described in LPB (Cai et al. 2014). Our results indicate a preferential innervation of LPB by SOM+, not by PKCδ neurons, in both STLD and CeL/C. Furthermore, we revealed by anterograde tracing that the axonal varicosities from EAc can specifically target CGRP, as well as non-CGRP neurons in LPBE.

Similarly, PAG-projecting neurons in STL and CeA have been known to express multiple neuronal markers such as neurotensin, CRF, and SOM (Gray and Magnuson 1992). In CeL/C, SOM+ neurons, but not PKCδ ones, have been shown to project to PAG by tract-tracing in SOM-cre mouse line (Penzo et al. 2014). So far, our findings on PAG/DR-projecting neurons are consistent with what has been reported for CeA and suggest that the same organization may exist in the STLD-PAG pathway. Beside LPB and PAG, CeL/C SOM+ neurons can also project to the solitary nucleus (Sol) (Higgins and Schwaber 1983; Gray and Magnuson 1987) and to the paraventricular thalamic nucleus (Penzo et al. 2014). Taken together, it is reasonable to hypothesize that SOM+ neurons, not PKCδ+ ones, are the major long-range projection neurons in STLD and CeL/C. However, SOM+ cells might not be the only population involved in long range projections. Indeed, several neuropeptidic markers, including ENK, CRF and neurotensin, have also been detected in brainstem-projecting neurons of CeL/C and STLD, but also of CeM and STLV (Gray and Magnuson 1992, 1987; Moga and Gray 1985; Moga et al. 1989; Magableh and Lundy 2014).

#### Functional implications of cell-type specific circuits in EAc

The pioneer studies of Cassell’s group (Cassell et al. 1986; Sun et al. 1991; Sun and Cassell 1993; Sun et al. 1994; Cassell et al. 1999) established the notion that in the rat CeA, CeL (and CeC) constitute an inhibitory interface between extra-EAc inputs and the CeA outputs derived from CeM. The organization of this microcircuitry was later precised in mice to show that, in fear conditioning, a conditioned stimulus, previously associated to an unconditioned stimulus, activates in CeL/C a population of PKCδ negative cells, potentially SOM+, which inhibits in turn a population of PKCδ+ cells projecting to CeM, leading thus to the disinhibition of the CeM outputs neurons (Ciocchi et al. 2010; Haubensak et al. 2010). Subsequent studies have detailed the roles of CeL/C PKCδ+ and SOM+ cells, along with LPB CGRP input, in fear learning and memory, in fear generalization and anxiety (Li et al. 2013; Han et al. 2015; Botta et al. 2015; Penzo et al. 2015). The role of these CeA circuits in feeding has also been examined through elegant studies showing that LPB CGRP signaling to PKCδ+ CeL/C suppresses appetite, while other inputs, including those from BL, can target other cell populations (i.e. SOM+ and Htr2a+) that promote appetite (Carter et al. 2013; Cai et al. 2014; Campos et al. 2016; Douglass et al. 2017; Kim et al. 2017). The CeA circuit we described here is consistent with the connectivity revealed in these studies. By contrast, this level of precision in microcircuits has not yet been reached for STL. The STL has been shown to be largely involved in contextual fear learning, anxiety and stress response (Zimmerman and Maren 2011; Goode et al. 2015; Daldrup et al. 2016; De Bundel et al. 2016; Davis et al. 2009). De Bundel and colleagues showed that fear generalization relays on a coordinated action of STLD and CeL/C dopamine D2 receptor-expressing neurons, which mostly coexpress PKCδ (De Bundel et al. 2016). Thus, considering the parallel circuits existing in CeL/C and STLD, it is possible that LPB ➜STLD pathway use a similar microcircuitry than CeL/C to support STL roles in associative learning and memory or in feeding.

### Conclusions

Although the principle components of EAc are well-known to substantially share input/output connectivities and neurochemical features, comparative studies of STL and CeA neuronal circuits at cellular level are missing. In this study, we revealed a new depth of structural similarity between STLD and CeL/C by showing the existence of similar cell-type specific neuronal circuits in both nuclei. We showed that, like in CeA, the non-overlapping PKCδ+ and SOM+ neuronal populations also exist in STLD. In both nuclei, these two distinct neuronal groups form cell-type specific microcircuits integrating long-range inputs, mediating intrinsic connections, and sending long-range projections. In addition, these parallel microcircuits are, at the same time, integrated circuits, largely through interconnections within nuclei, between STLD and CeL/C and from STLD to CeM as well as from CeL/C to STLV.

ST and CeA are also known to be similarly involved in emotion, but with distinct roles. For instance, both structures have been implicated in fear and anxiety, with ST being more involved in unconditioned/sustained fear response or anxiety-like behavior versus CeA being more implicated in conditioned/phasic fear response (Walker and Davis 1997; Walker et al. 2003; Davis et al. 2009; Lebow and Chen 2016). Similarly, the CeA participates in both sensory and affective aspects of pain (Neugebauer et al. 2004; Carrasquillo and Gereau 2007; Neugebauer 2015; Veinante et al. 2013), while ST seems to contribute mainly to the affective component of pain (Deyama et al. 2008; Minami and Ide 2015). So far, it is not clear what kind of structural differences underlies such functional discrepancy in ST and CeA. One possibility could be the existence of subtle differences in the inputs and outputs circuits of ST and CeA, as well as in the specificity of local neuronal pools. For example, it is remain to be explored whether ST and CeA are innervated by different sets of neurons in LPB or InsCx, or whether different pools of PKCδ+ or SOM+ neurons are involved in fear versus anxiety. Another explanation could be the asymmetric connections between STLD and CeA, where the projection from CeA to STL seems to be stronger than that of the reverse direction (Dong et al. 2001a; Oler et al. 2017). Again, the functional implications of these structural differences remain to be further explored.

So far, compared to STL, the structures and functions of CeA microcircuits have been better studied by cell-type/pathway specific genetic manipulation and behavioral assays (Ciocchi et al. 2010; Haubensak et al. 2010; Cai et al. 2014; Han et al. 2015; Li et al. 2013). Our results demonstrate that CeA-like microcircuits also exist in STL, and that they contribute to a complex network linking the components of the EAc. Future studies on structures and functions of neuronal circuits of ST might benefit from considering previous researches of CeA microcircuits.

## Acknowledgments

We thank Dr. Paul Klosen for helpful advices in CARD method and Dr. Alessandro Bilella for help in using NanoZoomer S60 platform. This work was supported by the Centre National de la Recherche Scientifique (contract UPR3212), the University of Strasbourg and the NeuroTime Erasmus Mundus Joint Doctorate Program.

## Compliance with ethical standards

All applicable international, national, and/or institutional guidelines for the care and use of animals were followed.

## Informed consent

No human subject were used in this study

## Disclosure of potential conflicts of interest

The authors declare that they have no conflict of interest.

## References

Alden M, Besson JM, Bernard JF (1994) Organization of the efferent projections from the pontine parabrachial area to the bed nucleus of the stria terminalis and neighboring regions: a PHA-L study in the rat. J Comp Neurol 341 (3):289–314. doi:10.1002/cne.903410302

Alheid GF (2003) Extended amygdala and basal forebrain. Ann N Y Acad Sci 985:185–205. doi: 10.1111/j.1749-6632.2003.tb07082.x

Bernard JF, Alden M, Besson JM (1993) The organization of the efferent projections from the pontine parabrachial area to the amygdaloid complex: a Phaseolus vulgaris leucoagglutinin (PHA-L) study in the rat. J Comp Neurol 329 (2):201–229. doi:10.1002/cne.903290205

Botta P, Demmou L, Kasugai Y, Markovic M, Xu C, Fadok JP, Lu T, Poe MM, Xu L, Cook JM, Rudolph U, Sah P, Ferraguti F, Luthi A (2015) Regulating anxiety with extrasynaptic inhibition. Nat Neurosci 18 (10): 1493–1500. doi:10.1038/nn.4102

Cai H, Haubensak W, Anthony TE, Anderson DJ (2014) Central amygdala PKC-delta(+) neurons mediate the influence of multiple anorexigenic signals. Nat Neurosci 17 (9):1240–1248. doi: 10.1038/nn.3767

Campos CA, Bowen AJ, Schwartz MW, Palmiter RD (2016) Parabrachial CGRP Neurons Control Meal Termination. Cell Metab 23 (5):811–820. doi:10.1016/j.cmet.2016.04.006

Carrasquillo Y, Gereau RWt (2007) Activation of the extracellular signal-regulated kinase in the amygdala modulates pain perception. J Neurosci 27 (7): 1543–1551. doi: 10.1523/JNEUROSCI.3536-06.2007

Carter ME, Soden ME, Zweifel LS, Palmiter RD (2013) Genetic identification of a neural circuit that suppresses appetite. Nature 503 (7474): 111–114. doi:10.1038/nature12596

Cassell MD, Freedman LJ, Shi C (1999) The intrinsic organization of the central extended amygdala. Ann N Y Acad Sci 877:217–241

Cassell MD, Gray TS, Kiss JZ (1986) Neuronal architecture in the rat central nucleus of the amygdala: a cytological, hodological, and immunocytochemical study. J Comp Neurol 246 (4):478–499. doi:10.1002/cne.902460406

Chieng BC, Christie MJ, Osborne PB (2006) Characterization of neurons in the rat central nucleus of the amygdala: cellular physiology, morphology, and opioid sensitivity. J Comp Neurol 497 (6):910–927. doi: 10.1002/cne.21025

Ciocchi S, Herry C, Grenier F, Wolff SB, Letzkus JJ, Vlachos I, Ehrlich I, Sprengel R, Deisseroth K, Stadler MB, Muller C, Luthi A (2010) Encoding of conditioned fear in central amygdala inhibitory circuits. Nature 468 (7321):277–282. doi:10.1038/nature09559

D’Hanis W, Linke R, Yilmazer-Hanke DM (2007) Topography of thalamic and parabrachial calcitonin gene-related peptide (CGRP) immunoreactive neurons projecting to subnuclei of the amygdala and extended amygdala. J Comp Neurol 505 (3):268–291. doi:10.1002/ene.21495

Daldrup T, Lesting J, Meuth P, Seidenbecher T, Pape HC (2016) Neuronal correlates of sustained fear in the anterolateral part of the bed nucleus of stria terminalis. Neurobiol Learn Mem 131:137–146. doi: 10.1016/j.nlm.2016.03.020

Davis M, Shi C (1999) The extended amygdala: are the central nucleus of the amygdala and the bed nucleus of the stria terminalis differentially involved in fear versus anxiety? Ann N Y Acad Sci 877:281–291

Davis M, Walker DL, Miles L, Grillon C (2009) Phasic vs sustained fear in rats and humans: role of the extended amygdala in fear vs anxiety. Neuropsychopharmacology 35 (1): 105–135. doi: 10.1038/npp.2009.109

De Bundel D, Zussy C, Espallergues J, Gerfen CR, Girault JA, Valjent E (2016) Dopamine D2 receptors gate generalization of conditioned threat responses through mTORC1 signaling in the extended amygdala. Mol Psychiatry 21 (11): 1545–1553. doi:10.1038/mp.2015.210

de Olmos JS, Heimer L (1999) The concepts of the ventral striatopallidal system and extended amygdala. Ann N Y Acad Sci 877:1–32

Delaney AJ, Crane JW, Sah P (2007) Noradrenaline modulates transmission at a central synapse by a presynaptic mechanism. Neuron 56 (5):880–892. doi:10.1016/j.neuron.2007.10.022

Deyama S, Katayama T, Ohno A, Nakagawa T, Kaneko S, Yamaguchi T, Yoshioka M, Minami M (2008) Activation of the beta-adrenoceptor-protein kinase A signaling pathway within the ventral bed nucleus of the stria terminalis mediates the negative affective component of pain in rats. J Neurosci 28 (31):7728–7736. doi:10.1523/JNEUROSCI.1480-08.2008

Dobolyi A, Irwin S, Makara G, Usdin TB, Palkovits M (2005) Calcitonin gene-related peptide-containing pathways in the rat forebrain. J Comp Neurol 489 (1):92–119. doi: 10.1002/cne.20618

Dong HW, Petrovich GD, Swanson LW (2001a) Topography of projections from amygdala to bed nuclei of the stria terminalis. Brain Res Brain Res Rev 38 (1-2): 192–246

Dong HW, Petrovich GD, Watts AG, Swanson LW (2001b) Basic organization of projections from the oval and fusiform nuclei of the bed nuclei of the stria terminalis in adult rat brain. J Comp Neurol 436 (4):430–455

Douglass AM, Kucukdereli H, Ponserre M, Markovic M, Grundemann J, Strobel C, Alcala Morales PL, Conzelmann KK, Luthi A, Klein R (2017) Central amygdala circuits modulate food consumption through a positive-valence mechanism. Nat Neurosci 20 (10): 1384–1394. doi:10.1038/nn.4623

Fadok JP, Krabbe S, Markovic M, Courtin J, Xu C, Massi L, Botta P, Bylund K, Muller C, Kovacevic A, Tovote P, Luthi A (2017) A competitive inhibitory circuit for selection of active and passive fear responses. Nature 542 (7639):96–100. doi:10.1038/nature21047

Goode TD, Kim JJ, Maren S (2015) Reversible Inactivation of the Bed Nucleus of the Stria Terminalis Prevents Reinstatement But Not Renewal of Extinguished Fear. eNeuro 2 (3). doi: 10.1523/ENEURO.0037-15.2015

Gray TS, Magnuson DJ (1987) Neuropeptide neuronal efferents from the bed nucleus of the stria terminalis and central amygdaloid nucleus to the dorsal vagal complex in the rat. J Comp Neurol 262 (3):365–374. doi:10.1002/cne.902620304

Gray TS, Magnuson DJ (1992) Peptide immunoreactive neurons in the amygdala and the bed nucleus of the stria terminalis project to the midbrain central gray in the rat. Peptides 13 (3):451–460

Gungor NZ, Pare D (2016) Functional Heterogeneity in the Bed Nucleus of the Stria Terminalis. J Neurosci 36 (31):8038–8049. doi:10.1523/JNEUROSCI.0856-16.2016

Gungor NZ, Yamamoto R, Pare D (2015) Optogenetic study of the projections from the bed nucleus of the stria terminalis to the central amygdala. Journal of Neurophysiology 114 (5):2903–2911. doi: 10.1152/jn.00677.2015

Han S, Soleiman MT, Soden ME, Zweifel LS, Palmiter RD (2015) Elucidating an Affective Pain Circuit that Creates a Threat Memory. Cell 162 (2):363–374. doi:10.1016/j.cell.2015.05.057

Haubensak W, Kunwar PS, Cai H, Ciocchi S, Wall NR, Ponnusamy R, Biag J, Dong HW, Deisseroth K, Callaway EM, Fanselow MS, Luthi A, Anderson DJ (2010) Genetic dissection of an amygdala microcircuit that gates conditioned fear. Nature 468 (7321):270–276. doi:10.1038/nature09553

Higgins GA, Schwaber JS (1983) Somatostatinergic projections from the central nucleus of the amygdala to the vagal nuclei. Peptides 4 (5):657–662

Hunt S, Sun Y, Kucukdereli H, Klein R, Sah P (2017) Intrinsic Circuits in the Lateral Central Amygdala. eNeuro 4 (1). doi:10.1523/ENEURO.0367-16.2017

Hunyady B, Krempels K, Harta G, Mezey E (1996) Immunohistochemical signal amplification by catalyzed reporter deposition and its application in double immunostaining. J Histochem Cytochem 44 (12): 1353–1362

Janak PH, Tye KM (2015) From circuits to behaviour in the amygdala. Nature 517 (7534):284–292. doi: 10.1038/nature14188

Jennings JH, Sparta DR, Stamatakis AM, Ung RL, Pleil KE, Kash TL, Stuber GD (2013) Distinct extended amygdala circuits for divergent motivational states. Nature 496 (7444):224–228. doi: 10.1038/nature12041

Jhou TC, Geisler S, Marinelli M, Degarmo BA, Zahm DS (2009) The mesopontine rostromedial tegmental nucleus: A structure targeted by the lateral habenula that projects to the ventral tegmental area of Tsai and substantia nigra compacta. J Comp Neurol 513 (6):566–596. doi: 10.1002/cne.21891

Kim J, Zhang X, Muralidhar S, LeBlanc SA, Tonegawa S (2017) Basolateral to Central Amygdala Neural Circuits for Appetitive Behaviors. Neuron 93 (6):1464–1479 e1465. doi: 10.1016/j.neuron.2017.02.034

Kim SY, Adhikari A, Lee SY, Marshel JH, Kim CK, Mallory CS, Lo M, Pak S, Mattis J, Lim BK, Malenka RC, Warden MR, Neve R, Tye KM, Deisseroth K (2013) Diverging neural pathways assemble a behavioural state from separable features in anxiety. Nature 496 (7444):219–223. doi: 10.1038/nature12018

Lebow MA, Chen A (2016) Overshadowed by the amygdala: the bed nucleus of the stria terminalis emerges as key to psychiatric disorders. Mol Psychiatry 21 (4):450–463. doi:10.1038/mp.2016.1

Lein ES, et al. (2007) Genome-wide atlas of gene expression in the adult mouse brain. Nature 445 (7124): 168–176. doi:10.1038/nature05453

Li H, Penzo MA, Taniguchi H, Kopec CD, Huang ZJ, Li B (2013) Experience-dependent modification of a central amygdala fear circuit. Nat Neurosci 16 (3):332–339. doi: 10.1038/nn.3322

Magableh A, Lundy R (2014) Somatostatin and corticotrophin releasing hormone cell types are a major source of descending input from the forebrain to the parabrachial nucleus in mice. Chem Senses 39 (8):673–682. doi:10.1093/chemse/bju038

Mazzone CM, Pati D, Michaelides M, DiBerto J, Fox JH, Tipton G, Anderson C, Duffy K, McKlveen JM, Hardaway JA, Magness ST, Falls WA, Hammack SE, McElligott ZA, Hurd YL, Kash TL (2016) Acute engagement of Gq-mediated signaling in the bed nucleus of the stria terminalis induces anxiety-like behavior. Mol Psychiatry. doi:10.1038/mp.2016.218

McDonald AJ (1982) Cytoarchitecture of the central amygdaloid nucleus of the rat. J Comp Neurol 208 (4):401–418. doi:10.1002/cne.902080409

McDonald AJ (1991) Topographical organization of amygdaloid projections to the caudatoputamen, nucleus accumbens, and related striatal-like areas of the rat brain. Neuroscience 44 (1): 15–33

McDonald AJ, Shammah-Lagnado SJ, Shi C, Davis M (1999) Cortical afferents to the extended amygdala. Ann N Y Acad Sci 877:309–338

Minami M, Ide S (2015) How does pain induce negative emotion? Role of the bed nucleus of the stria terminalis in pain-induced place aversion. Curr Mol Med 15 (2): 184–190

Moga MM, Gray TS (1985) Evidence for corticotropin-releasing factor, neurotensin, and somatostatin in the neural pathway from the central nucleus of the amygdala to the parabrachial nucleus. J Comp Neurol 241 (3):275–284. doi:10.1002/cne.902410304

Moga MM, Saper CB, Gray TS (1989) Bed nucleus of the stria terminalis: cytoarchitecture, immunohistochemistry, and projection to the parabrachial nucleus in the rat. J Comp Neurol 283 (3):315–332. doi: 10.1002/cne.902830302

Neugebauer V (2015) Amygdala pain mechanisms. Handb Exp Pharmacol 227:261–284. doi: 10.1007/978-3-662-46450-2_13

Neugebauer V, Li W, Bird GC, Han JS (2004) The amygdala and persistent pain. Neuroscientist 10 (3):221–234. doi: 10.1177/1073858403261077

Oh SW, Harris JA, Ng L, Winslow B, Cain N, Mihalas S, Wang Q, Lau C, Kuan L, Henry AM, Mortrud MT, Ouellette B, Nguyen TN, Sorensen SA, Slaughterbeck CR, Wakeman W, Li Y, Feng D, Ho A, Nicholas E, Hirokawa KE, Bohn P, Joines KM, Peng H, Hawrylycz MJ, Phillips JW, Hohmann JG, Wohnoutka P, Gerfen CR, Koch C, Bernard A, Dang C, Jones AR, Zeng H (2014) A mesoscale connectome of the mouse brain. Nature 508 (7495):207–214. doi:10.1038/nature13186

Oler JA, Tromp DP, Fox AS, Kovner R, Davidson RJ, Alexander AL, McFarlin DR, Birn RM, B EB, deCampo DM, Kalin NH, Fudge JL (2017) Connectivity between the central nucleus of the amygdala and the bed nucleus of the stria terminalis in the non-human primate: neuronal tract tracing and developmental neuroimaging studies. Brain Struct Funct 222 (1):21–39. doi:10.1007/s00429-016-1198-9

Panguluri S, Saggu S, Lundy R (2009) Comparison of somatostatin and corticotrophin-releasing hormone immunoreactivity in forebrain neurons projecting to taste-responsive and non-responsive regions of the parabrachial nucleus in rat. Brain Res 1298:57–69. doi:10.1016/j.brainres.2009.08.038

Paxinos G, Franklin K (2012) Paxinos and Franklin’s The mouse brain in stereotaxic coordinates. Fourth edition. edn. Academic Press, Amsterdam

Penzo MA, Robert V, Li B (2014) Fear conditioning potentiates synaptic transmission onto long-range projection neurons in the lateral subdivision of central amygdala. J Neurosci 34 (7):2432–2437. doi: 10.1523/JNEUROSCI.4166-13.2014

Penzo MA, Robert V, Tucciarone J, De Bundel D, Wang M, Van Aelst L, Darvas M, Parada LF, Palmiter RD, He M, Huang ZJ, Li B (2015) The paraventricular thalamus controls a central amygdala fear circuit. Nature. doi:10.1038/nature13978

Petrovich GD, Swanson LW (1997) Projections from the lateral part of the central amygdalar nucleus to the postulated fear conditioning circuit. Brain Res 763 (2):247–254

Pitkanen A, Savander M, Nurminen N, Ylinen A (2003) Intrinsic synaptic circuitry of the amygdala. Ann N Y Acad Sci 985:34–49

Pitkanen A, Stefanacci L, Farb CR, Go GG, LeDoux JE, Amaral DG (1995) Intrinsic connections of the rat amygdaloid complex: projections originating in the lateral nucleus. J Comp Neurol 356 (2):288–310. doi: 10.1002/cne.903560211

Pomrenze MB, Millan EZ, Hopf FW, Keiflin R, Maiya R, Blasio A, Dadgar J, Kharazia V, De Guglielmo G, Crawford E, Janak PH, George O, Rice KC, Messing RO (2015) A Transgenic Rat for Investigating the Anatomy and Function of Corticotrophin Releasing Factor Circuits. Front Neurosci 9:487. doi:10.3389/fnins.2015.00487

Salio C, Averill S, Priestley JV, Merighi A (2007) Costorage of BDNF and neuropeptides within individual dense-core vesicles in central and peripheral neurons. Dev Neurobiol 67 (3):326–338. doi:10.1002/dneu.20358

Saper CB (1982) Convergence of autonomic and limbic connections in the insular cortex of the rat. J Comp Neurol 210 (2):163–173. doi:10.1002/cne.902100207

Sarhan M, Freund-Mercier MJ, Veinante P (2005) Branching patterns of parabrachial neurons projecting to the central extended amgydala: single axonal reconstructions. J Comp Neurol 491 (4):418–442. doi: 10.1002/cne.20697

Sato M, Ito M, Nagase M, Sugimura YK, Takahashi Y, Watabe AM, Kato F (2015) The lateral parabrachial nucleus is actively involved in the acquisition of fear memory in mice. Mol Brain 8:22. doi:10.1186/s13041-015-0108-z

Savander V, Go CG, Ledoux JE, Pitkanen A (1996) Intrinsic connections of the rat amygdaloid complex: projections originating in the accessory basal nucleus. J Comp Neurol 374 (2):291–313. doi: 10.1002/(SICI)1096-9861(19961014)374:2…lt;291::AID-CNE10…gt;3.0.CO;2-Y

Schindelin J, Arganda-Carreras I, Frise E, Kaynig V, Longair M, Pietzsch T, Preibisch S, Rueden C, Saalfeld S, Schmid B, Tinevez JY, White DJ, Hartenstein V, Eliceiri K, Tomancak P, Cardona A (2012) Fiji: an open-source platform for biological-image analysis. Nat Methods 9 (7):676–682. doi: 10.1038/nmeth.2019

Shackman AJ, Fox AS (2016) Contributions of the Central Extended Amygdala to Fear and Anxiety. J Neurosci 36 (31):8050–8063. doi:10.1523/JNEUROSCI.0982-16.2016

Speel EJ, Ramaekers FC, Hopman AH (1997) Sensitive multicolor fluorescence in situ hybridization using catalyzed reporter deposition (CARD) amplification. J Histochem Cytochem 45 (10): 1439–1446

Sun N, Cassell MD (1993) Intrinsic GABAergic neurons in the rat central extended amygdala. J Comp Neurol 330 (3):381–404. doi:10.1002/cne.903300308

Sun N, Roberts L, Cassell MD (1991) Rat central amygdaloid nucleus projections to the bed nucleus of the stria terminalis. Brain Res Bull 27 (5):651–662

Sun N, Yi H, Cassell MD (1994) Evidence for a GABAergic interface between cortical afferents and brainstem projection neurons in the rat central extended amygdala. J Comp Neurol 340 (1):43–64. doi: 10.1002/cne.903400105

Tokita K, Inoue T, Boughter JD, Jr. (2009) Afferent connections of the parabrachial nucleus in C57BL/6J mice. Neuroscience 161 (2):475–488. doi:10.1016/j.neuroscience.2009.03.046

Veinante P, Freund-Mercier MJ (1998) Intrinsic and extrinsic connections of the rat central extended amygdala: an in vivo electrophysiological study of the central amygdaloid nucleus. Brain Res 794 (2): 188–198. doi:S0006-8993(98)00228-5 [pii]

Veinante P, Freund-Mercier MJ (2003) Branching Patterns of Central Amygdaloid Nucleus Efferents in the Rat: Single-Axon Reconstructions. Ann N Y Acad Sci 985:552–553. doi: 10.1111/j.749-6632.2003.tb07126.x

Veinante P, Stoeckel ME, Freund-Mercier MJ (1997) GABA- and peptide-immunoreactivities co-localize in the rat central extended amygdala. Neuroreport 8 (13):2985–2989

Veinante P, Yalcin I, Barrot M (2013) The amygdala between sensation and affect: a role in pain. J Mol Psychiatry 1 (1):9. doi:10.1186/2049-9256-1-9

Walker DL, Davis M (1997) Double dissociation between the involvement of the bed nucleus of the stria terminalis and the central nucleus of the amygdala in startle increases produced by conditioned versus unconditioned fear. J Neurosci 17 (23):9375–9383

Walker DL, Toufexis DJ, Davis M (2003) Role of the bed nucleus of the stria terminalis versus the amygdala in fear, stress, and anxiety. Eur J Pharmacol 463 (1-3): 199–216. doi:S0014299903012822 [pii]

Weller KL, Smith DA (1982) Afferent connections to the bed nucleus of the stria terminalis. Brain Res 232 (2):255–270

Wood J, Verma D, Lach G, Bonaventure P, Herzog H, Sperk G, Tasan RO (2016) Structure and function of the amygdaloid NPY system: NPY Y2 receptors regulate excitatory and inhibitory synaptic transmission in the centromedial amygdala. Brain Struct Funct 221 (7):3373–3391. doi: 10.1007/s00429-015-1107-7

Yasui Y, Breder CD, Saper CB, Cechetto DF (1991a) Autonomic responses and efferent pathways from the insular cortex in the rat. J Comp Neurol 303 (3):355–374. doi:10.1002/cne.903030303

Yasui Y, Saper CB, Cechetto DF (1991b) Calcitonin gene-related peptide (CGRP) immunoreactive projections from the thalamus to the striatum and amygdala in the rat. J Comp Neurol 308 (2):293–310. doi: 10.1002/cne.903080212

Yu K, Garcia da Silva P, Albeanu DF, Li B (2016) Central Amygdala Somatostatin Neurons Gate Passive and Active Defensive Behaviors. J Neurosci 36 (24):6488–6496. doi:10.1523/JNEUROSCI.4419-15.2016

Zimmerman JM, Maren S (2011) The bed nucleus of the stria terminalis is required for the expression of contextual but not auditory freezing in rats with basolateral amygdala lesions. Neurobiol Learn Mem 95 (2):199–205. doi:10.1016/j.nlm.2010.11.002

